# The murine lung microbiome is dysbalanced by the human pathogenic fungus *Aspergillus fumigatus* resulting in enrichment of anaerobic bacteria

**DOI:** 10.1101/2024.06.12.598217

**Authors:** Liubov Nikitashina, Xiuqiang Chen, Lukas Radosa, Kexin Li, Maria Straßburger, Bastian Seelbinder, Wibke Böhnke, Sarah Vielreicher, Sandor Nietzsche, Thorsten Heinekamp, Ilse D. Jacobsen, Gianni Panagiotou, Axel A. Brakhage

## Abstract

Here, we report significant changes in the composition of the lung microbiome and metabolome of mice under immune suppression, infection of immunosuppressed mice with virulent and avirulent strains of the clinically important human-pathogenic fungus *Aspergillus fumigatus*, and treatment with the clinically used antifungal drug voriconazole. Our data also indicate the important role of the gut microbiome for the lung homeostasis mediated by the plasma metabolome. In the lung microbiome, infection by *A. fumigatus* led to a significant increase of anaerobic bacteria, most prominently of *Ligilactobacillus murinus,* which was confirmed by the isolation of live bacteria from the murine lower respiratory tract. *In vitro*, *L. murinus* is tolerated and even internalized by alveolar epithelial cells. Co-cultivation of *L. murinus* and *A. fumigatus* led to a reduction in oxygen concentration accompanied by an increase of *L. murinus* cells suggesting that *A. fumigatus* establishes a microaerophilic niche, thereby promoting growth of anaerobic bacteria.

## Introduction

For many years, the healthy lung was considered a sterile environment ^1^. This assumption has been challenged, however, when advanced molecular techniques, such as next-generation sequencing, allowed detection of DNA of various microorganisms in the lower respiratory tract. It was found that the load of microbial DNA in the lower respiratory tract is remarkably low compared to DNA detected in the upper respiratory tract and other host sites, such as the gut ^1,2^. This low amount of DNA may be due to the fact that although the lower respiratory tract is constantly exposed to living microorganisms by breathing, several mechanisms provide rapid clearance of microbes including mucociliary clearance and the activity of both, the innate and the adaptive immune system ^2,3^.

It is not surprising that the composition of the microbiome of the healthy lung considerably differs from that of the microbiome of a diseased lung ^1^. For example, the lung microbiome of patients with inflammatory respiratory diseases such as cystic fibrosis, chronic obstructive pulmonary disease, and idiopathic pulmonary fibrosis usually has a lower bacterial diversity and the apparent dysbiosis of the lung microbiome is associated with a worsening of the course of the disease ^4–8^. Furthermore, features of lung microbial dysbiosis were shown to correlate with and to be predictive of the clinical outcome of patients with sepsis, acute respiratory distress syndrome, and invasive aspergillosis ^9,10^.

An important aspect of studies on the lung microbiome concerns the identification of resident microbiota in a healthy lung and how the microbiota changes during dysbiosis caused by immune suppression, infection, and therapy. It is still a matter of debate how the alteration of metabolite composition during dysbiosis affects the gut-lung axis. Finally, the major challenge is the identification of cause-effect-relationships, *i.e*., the elucidation of causes of changes of the microbiome under dysbiosis conditions and how these changes influence the outcome of an infection.

To address these questions, we performed an in-depth analysis of the lung microbiome of laboratory specific, pathogen-free (SPF) mice before and after infection with the clinically relevant fungus *Aspergillus fumigatus*. *A. fumigatus* is the most important air-borne fungal pathogen causing life-threatening invasive aspergillosis ^10–14^.

To investigate the influence of invasive aspergillosis not only on the lung but also on the gut-lung axis, we analyzed the gut microbiome, and the metabolome of both the plasma and the lung. In addition, immunocompromised mice were infected with conidia of either a virulent wild-type or an avirulent mutant strain of *A. fumigatus*. This enabled us to determine specific interactions during infection between the pathogen and the lung microbiota. Finally, isolation of culturable microorganisms from the mouse lower respiratory tract allowed us to obtain insight into dependencies of the pathogen and members of the lung microbiomes.

## Results

### *Ligilactobacillus* dominates the murine lung microbiome

To investigate the response of the lung microbiome to medical interventions and pulmonary infection with *A. fumigatus*, we used a mouse model for invasive aspergillosis. We analyzed the lung microbiome and metabolome, plasma metabolome and gut microbiome of healthy mice and mice during immunosuppression caused by cortisone acetate treatment, because *A. fumigatus* mainly infects immunocompromised human individuals ^11^. We also investigated the systemic effect of antifungal treatment with the clinically used antifungal drug voriconazole on the microbiomes and metabolomes (Figure 1A).

**Figure 1:**
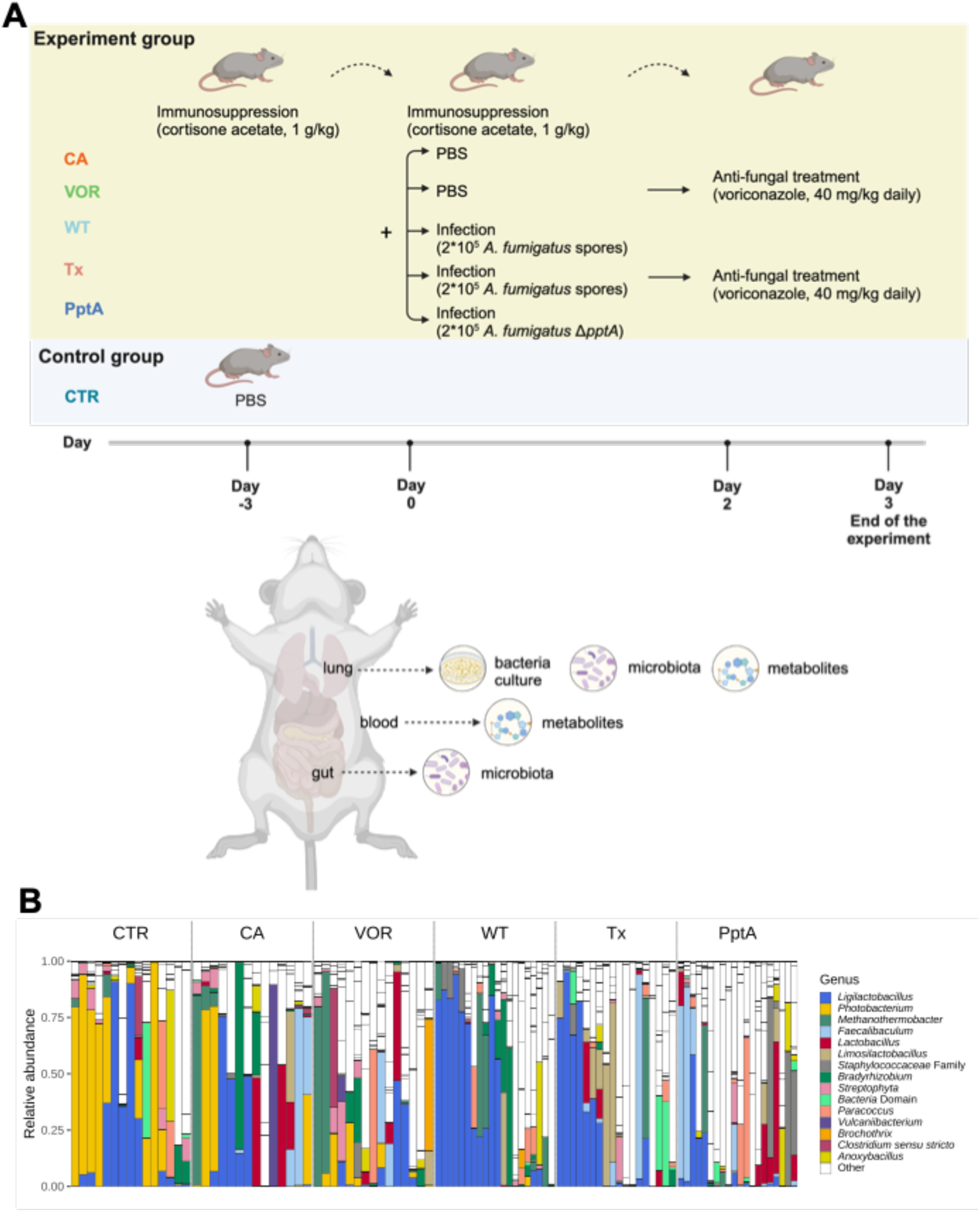
Immunosuppression, *A. fumigatus* infection, and antifungal treatment have different impacts on the composition of the lung microbiome. A) Experimental design and illustration of a mouse with indication from which organs samples were taken for the different analyses. The graphic was created with BioRender. B) Composition of the lung microbiome of mice. Composition plot based on 16S rDNA. Abbreviations: control mice (CTR); cortisone acetate-immunosuppressed mice (CA); cortisone acetate-immunosuppressed mice treated with voriconazole for 3 days (VOR); cortisone acetate-immunosuppressed mice infected with *A. fumigatus* wild-type conidia (WT); mice immunosuppressed with cortisone acetate, infected with *A. fumigatus* and treated with voriconazole (Tx); and mice immunosuppressed with cortisone acetate and infected with conidia of the avirulent *A. fumigatus* Δ*pptA* strain (PptA). Each bar represents data from a single animal.

The composition of the lung microbiome was analyzed by 16S rDNA sequencing. As described previously ^15^, the lung bacterial composition was highly variable between the samples within each treatment group. The bacterial species whose DNA was detected in large quantities in the majority (91 of 102) of samples was identified as *Ligilactobacillus* (formerly named *Lactobacillus*) ^16^ (Figure 1B). In addition to *Ligilactobacillus*, *Photobacterium*, *Methanothermobacter*, *Faecalibaculum*, and *Lactobacillus* were the most abundant genera in the lung microbiome.

### Immunosuppression and antifungal treatment lead to alterations in the lung microbiome, gut microbiome, and metabolome

#### Lung and gut microbiome

Since in general immunosuppression is required for the development of invasive aspergillosis, mice were immunosuppressed with cortisone acetate (Figure 1A). To investigate the influence of antifungal treatment, immunosuppressed mice were treated daily with voriconazole. Since immunosuppression and antifungal treatment may have a substantial impact by themselves, we first analyzed changes in the lung microbiome, gut microbiome, lung metabolome and plasma metabolome without *A. fumigatus* infection.

We observed no significant differences in the alpha diversity of the lung microbiome between immunosuppressed and control mice (Figure 2A). By contrast, the gut microbiome richness of immunosuppressed mice was significantly increased (Chao index, *p* = 3.6 × 10^-5^, Wilcoxon rank-sum test), and entropy significantly decreased (Shannon index, *p* = 0.046, Wilcoxon rank-sum test) compared to the control mice (Figure 2B). For the lung microbiome, beta diversity was not significantly different between control and immunosuppressed mice (weighted UniFrac, *p* > 0.05, PERMANOVA). However, we observed a significant difference in the abundance of 8 genera (*p* < 0.05, |log_2_FC| > log_2_(1.5), DESeq2), with the largest change seen for *Bifidobacterium* (log_2_FC = 22.695, *p* = 1.2 × 10^-14^, FDR = 5 × 10^-13^, DESeq2) (Figure 2A, Table S1). This was different from the gut microbiome, where we observed a significant difference in beta diversity between control and immunosuppressed mice (*p* = 0.001, PERMANOVA). Under immunosuppression, the relative abundances of 23 genera changed significantly (*p* < 0.05, |log_2_FC| > log_2_(1.5), DESeq2), with the greatest change observed for *Turicibacter* (log_2_FC = 9.492, *p* = 4 × 10^-16^, FDR = 4.1 × 10^-15^) (Figure 2B, Table S2) whose strains are prominent members of the mammalian gut microbiota ^17^. It was recently shown that *T. sanguinis* has an important role in conferring resistance to mice against infections by *Citrobacter rodentium* ^18^.

**Figure 2:**
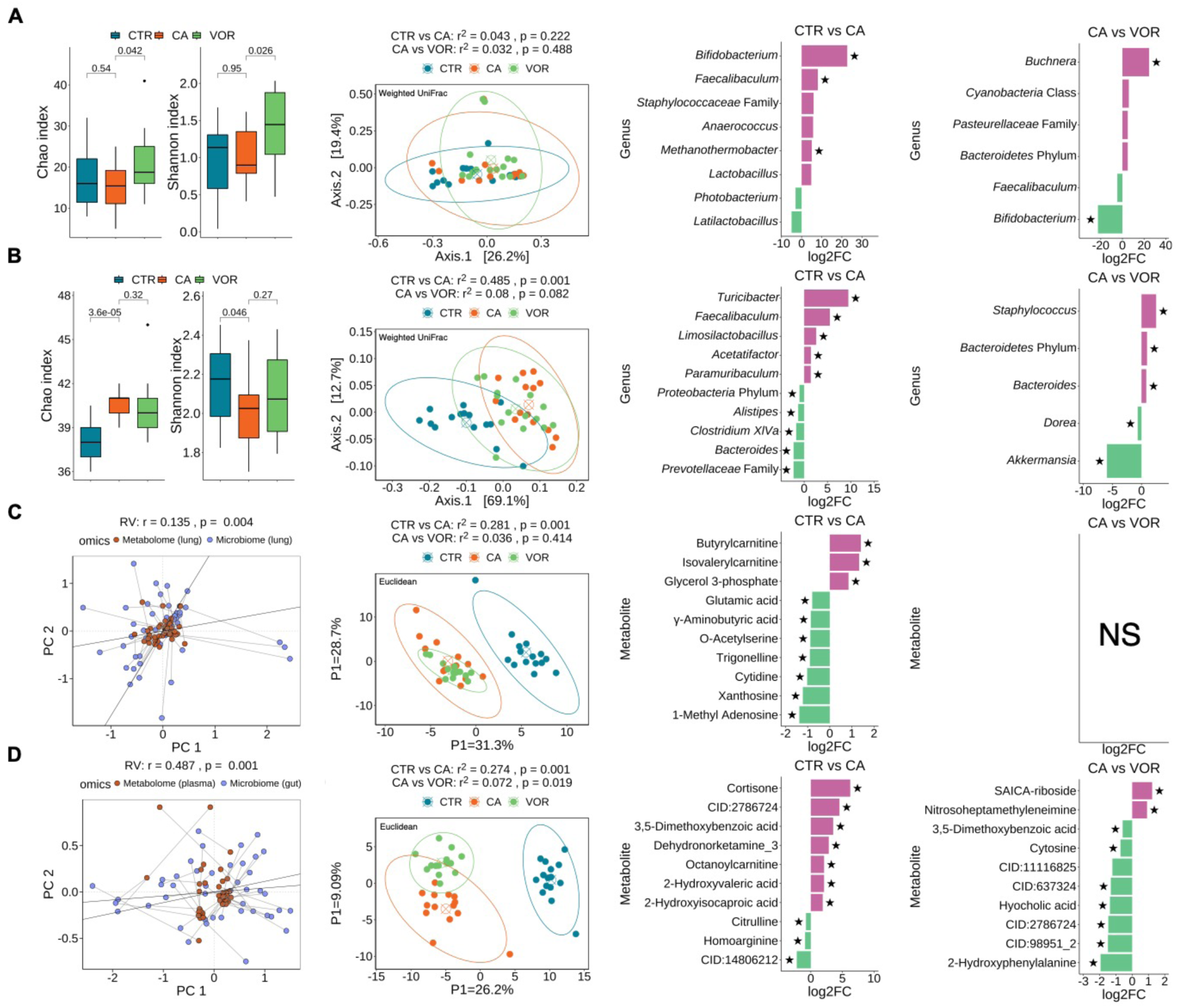
Immunosuppression and voriconazole treatment affect the lung microbiome, gut microbiome, lung metabolome and plasma metabolome. Control mice (CTR) were compared to mice immunosuppressed with cortisone acetate (CA). Immunosuppressed mice (CA) were compared to immunosuppressed mice treated with voriconazole (VOR). For A) the lung microbiome and B) the gut microbiome, alpha diversity was estimated using both the Chao index and the Shannon index. Two-sided Wilcoxon rank-sum test was used to assess the statistical significance. Beta diversity was estimated with the weighted UniFrac distance. PERMANOVA was used to assess the statistical significance. Genera with differential abundance (by *p* < 0.05 and |log_2_FC| > log_2_(1.5); DESeq2) are shown. When identification at the genus level was not possible, lower levels of identification to family or phylum level are indicated. Genera with FDR < 0.1 are marked with a black star. C) Correlation between lung microbiome and lung metabolome. The significance of RV values was estimated using permutation. The similarity between lung metabolome samples was determined using PLS-DA. PERMANOVA was used to assess the statistical significance. Metabolites with differential abundance (*p* < 0.05 and |log_2_FC| > log_2_(1.5); Wilcoxon rank-sum test) are shown. Metabolites with FDR < 0.1 are marked with a black star. D) Correlation between the gut microbiome and the plasma metabolome. The significance of RV values was estimated using permutation. The similarity between the plasma metabolome samples was determined using PLS-DA. PERMANOVA was used to assess the statistical significance. Metabolites with *p* < 0.05 and |log_2_FC| > log_2_(1.5) are regarded as differentially abundant; metabolites with FDR < 0.1 are indicated with a black star. Abbreviations: CID:2786724: 5,6-Dimethyl-4-oxo-4H-pyran-2-carboxylic acid; CID:14806212: 1,4a-Dimethyl-6-methylene-5-[2-(2-oxo-2,5-dihydro-3-furanyl)ethyl]decahydro-1-naphthalenecarboxylic acid; CID:11116825: 2-[(1S)-1-Hydroxyethyl]-4(1H)-quinazolinone; CID:637324: (8aR,12S,12aR)-12-Hydroxy-4-methyl-4,5,6,7,8,8a,12,12a-octahydro-2H-3-benzoxecine-2,9(1H)-dione; CID:2786724: 5,6-Dimethyl-4-oxo-4H-pyran-2-carboxylic acid; CID:98951: 3-(propan-2-yl)-octahydropyrrolo[1,2-a]pyrazine-1,4-dione. For differently abundant genera and metabolites, 10 genera or metabolites with the highest statistical significance are shown.

Voriconazole treatment had no significant effect on the overall composition of the lung microbiome (weighted UniFrac, *p* > 0.05, PERMANOVA). However, both community richness (Chao index, *p* = 0.042, Wilcoxon rank-sum test) and community entropy (Shannon index, *p* = 0.026, Wilcoxon rank-sum test) significantly increased. Furthermore, the relative abundances of 6 genera changed significantly (*p* < 0.05, |log_2_FC| > log_2_(1.5), DESeq2) (Figure 2A, Table S1). The effect of voriconazole treatment on the gut microbiome was negligible (Figure 2B, Table S2). The relative abundances of only 5 genera were statistically significantly altered (*p* < 0.05, |log_2_FC| > log_2_(1.5), DESeq2).

#### Lung and plasma metabolome

The gut microbiota has previously been shown to influence the composition of the plasma metabolome, particularly because metabolites entered the bloodstream ^19^. Here, we observed a moderate correlation between the gut microbiome and the plasma metabolome (RV, r = 0.487, *p* = 0.001, permutation test) and a lower, but also statistically significant, correlation between the lung microbiome and the lung metabolome (RV, r = 0.135, *p* = 0.004, permutation test) (Figure 2C,D). PLS-DA analysis revealed that both the lung and plasma metabolomes of immunocompromised mice were significantly different from those of control mice (Euclidean, *p* = 0.001, PERMANOVA). A significant difference was also observed in plasma samples (Euclidean, *p* = 0.019, PERMANOVA), but not in lung samples (*p* = 0.414, PERMANOVA), between immunocompromised mice and voriconazole-treated immunocompromised mice (Figure 2C,D). Among the metabolites that were statistically significantly altered by immunosuppression (*p* < 0.05, |log_2_FC| > log_2_(1.5), Wilcoxon rank-sum test), 8 metabolites were up-regulated and 16 were down-regulated in the lung samples (Figure 2C, Table S3), and 52 metabolites were up-regulated and 56 were down-regulated in the plasma samples (Figure 2D, Table S4). Of the increased plasma metabolites, bile acids and medium- and short-chain acylcarnitines correlated with the observed increase of *Turicibacter* in the gut microbiome. Strains of this genus are known to be able of bile acid deconjugation, leading to release of unconjugated bile acids, and modification of lipid metabolism ^17^. Along with this, butyrylcarnitine, isovalerylcarnitine, and propionylcarnitine were increased in the lung metabolome. Among the metabolites that changed statistically significantly during voriconazole treatment (*p* < 0.05, |log_2_FC| > log_2_(1.5), Wilcoxon rank-sum test), 3 metabolites were up-regulated and 11 were down-regulated in the plasma samples (Figure 2D, Table S4), but we found no differentially abundant metabolites in the lung.

We also observed a decrease of gamma-aminobutyric levels in the lungs upon immunosuppression. This might be connected with reduced functioning of immune system because gamma-aminobutyric acid was reported before to be produced by immune cells ^20^. In summary, we observe that immunosuppression with cortisone acetate had a major influence on the microbiomes and metabolomes irrespective of treatment with the antifungal voriconazole.

### Influence of *A. fumigatus* infection on microbiomes and metabolomes

To investigate the impact of *A. fumigatus* on microbiomes, we infected immunocompromised mice with conidia of the *A. fumigatus* wild-type strain and the Δ*pptA* mutant strain. The Δ*pptA* strain lacks the phosphopantetheinyl transferase, which is critical for the production of natural products, including virulence factors such as gliotoxin, the siderophores triacetylfusarinin C (TAFC) and ferricrocin, and dihydroxynaphthalene melanin ^21^. The Δ*pptA* mutant strain is therefore avirulent and cannot grow in the lung of immunosuppressed mice ^21^.

Samples were taken on day 3 after infection, at which immunocompromised mice infected with *A. fumigatus* wild type developed symptoms expected at this infection stage. Mice from other treatment groups did not show infection symptoms except of two immunocompromised infected with *A. fumigatus* wild type and treated with voriconazole mice, for which, however no signs of lung infection were observed (Figure S2, Table S5). We did not observe significant differences in beta-diversity of the lung microbiome after infection with either strain of *A. fumigatus* (Figure 3A, weighted UniFrac, *p* > 0.05, PERMANOVA). Interestingly, the gut microbiome indicated significant differences in immunocompromised mice compared to those infected with the *A. fumigatus* wild-type strain (weighted UniFrac, *p* = 0.001, PERMANOVA), but not with the avirulent *ΔpptA* strain (weighted UniFrac metric, *p* = 0.294, PERMANOVA). Significant differences between all three groups of mice were detected for both the lung and plasma metabolomes (Euclidean, *p* = 0.001, PERMANOVA) (Figure 3A).

**Figure 3:**
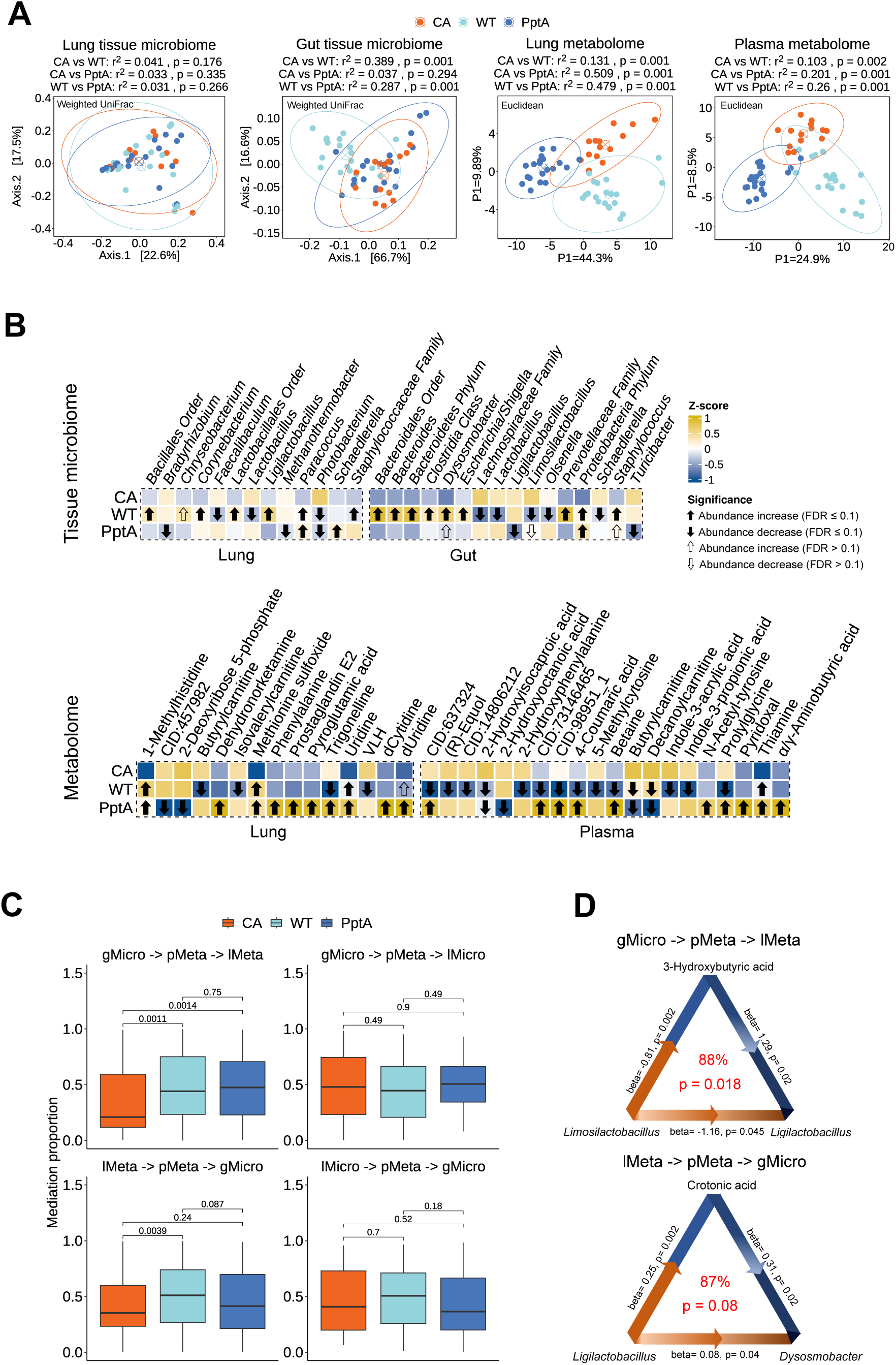
Infection with wild-type and avirulent Δ*pptA* conidia of *A. fumigatus* causes different effects on the lung microbiome, gut microbiome, lung metabolome and plasma metabolome (3 days post infection). Immunosuppressed mice (CA) were compared with immunosuppressed mice infected with *A. fumigatus* wild type (WT) or Δ*pptA* (PptA). A) Beta diversity [weighted UniFrac] of the lung and gut microbiome. PLS-DA for the lung and plasma metabolome. B) Heatmaps of differentially abundant genera (*p* < 0.05, |log_2_FC| > log_2_(1.5)). Z-transformed mean-abundances are shown. C) Mediation analysis of differentially abundant (*p* < 0.05, the union of CA *vs.* WT and CA *vs.* PptA) genera and metabolites in response to infection by the HIMA method. Top-Left: gut microbes (gMicro) influence lung metabolites (lMeta) through plasma metabolites (pMeta). Top-Right: gut microbes influence lung microbes (lMicro) through plasma metabolites. Bottom-Left: Lung metabolites influence gut microbes through plasma metabolites. Bottom-Right: Lung microbes influence gut microbes through plasma metabolites. D) Mediation analysis of lung bacterium *Ligilactobacillus*. Abbreviations: CID:457982: 10-(Hydroxymethyl)-4-methyl-8,11-dioxa-2,6-diazatricyclo[7.2.1.02,7]dodeca-3,6-dien-5-one; CID:637324: (8aR,12S,12aR)-12-Hydroxy-4-methyl-4,5,6,7,8,8a,12,12a-octahydro-2H-3-benzoxecine-2,9(1H)-dione; CID:14806212: 1,4a-Dimethyl-6-methylene-5-[2-(2-oxo-2,5-dihydro-3-furanyl)ethyl]decahydro-1-naphthalenecarboxylic acid; CID:73146465: 3-(1-hydroxyethyl)-2,3,6,7,8,8a-hexahydropyrrolo[1,2-a]pyrazine-1,4-dione; CID:98951: 3-(Propan-2-yl)-octahydropyrrolo[1,2-a]pyrazine-1,4-dione.

When comparing the abundance of bacterial genera between immunocompromised mice and immunocompromised mice infected with wild-type *A. fumigatus*, we observed that the relative abundance of 10 genera significantly changed in the lung microbiome. Among them, 4 genera are composed of obligate anaerobic species (*Ligilactobacillus*, *Lactobacillus*, *Faecalibaculum*, and a genus from the *Lactobacillales* order) and 4 genera of facultative anaerobic species (*Corynebacterium*, *Photobacterium*, and genera from *Staphylococcaceae* family and *Bacillales* order) (*p* < 0.05, |log_2_FC| > log_2_(1.5), DESeq2). For mice infected with *A. fumigatus* Δ*pptA*, the abundance of 5 genera was significantly altered, with 2 genera represented by anaerobic species, 1 genus by facultative anaerobic species, and 2 genera by aerobic species (Figure 3B, Table S1). Notably, lung samples infected with *A. fumigatus* wild-type, but not the avirulent *ΔpptA* strain, showed a significant increase in the relative abundance of the anaerobic *Ligilactobacillus* (*p* = 0.023, FDR = 0.097, log_2_FC = 2.835, DESeq2), the dominant species in all samples.

In the gut microbiome, the relative abundances of 13 genera were significantly altered by infection with *A. fumigatus* wild type (*p* < 0.05, |log_2_FC| > log_2_(1.5), DESeq2), with the highest change for *Escherichia/Shigella* whose relative abundance increased 20-fold (Figure 3B, Table S2). *Escherichia/Shigella* are considered to be facultative pathogens and frequently increase in abundance of these bacteria is a sign of a dysbiosis and indicates reduced immune protection ^22,23^. By contrast, infection with *A. fumigatus ΔpptA* only led to a significant change in the abundances of 6 genera (*p* < 0.05, |log_2_FC| > log_2_(1.5), DESeq2), including 3 genera with FDR less than 0.1 (black arrows). *Dysosmobacter* and *Staphylococcus* significantly increased and *Limosilactobacillus* significantly decreased in relative abundance in the gut of mice infected with both strains of *A. fumigatus*.

Among the metabolites with significantly altered abundance (*p* < 0.05, |log_2_FC| > log_2_(1.5), Wilcoxon rank-sum test) in the plasma samples, the majority of metabolites were decreased after infection with wild-type *A. fumigatus* (44 out of 48 significant metabolites), and only the abundance of thiamine, aspartic acid, creatine and 3-methylhistidine were increased. Infection with the avirulent strain of *A. fumigatus* resulted in an increase in the abundance of 56 metabolites and a decrease of 34 metabolites. In the plasma metabolome, among other metabolites, both infections caused a decrease of bile acids and tryptophan derivatives. The decrease of bile acids is potentially due to the decrease of *Turicibacter* in the gut microbiome, which was, however, not statistically significant for the *A. fumigatus* wild-type infection (*p* = 0.4).

In the lung metabolome, infection with the *A. fumigatus* wild type led to a significant increase in methionine sulfoxide, uridine and 1-methylhistidine, and a decrease in butyrylcarnitine, isovalerylcarnitine, trigonelline (nicotinic acid N-methylbetaine) and a putatively identified (Level 2b, see Materials and methods) tripeptide histidyl-leucyl-valine (*p* < 0.05, |log_2_FC| > log_2_(1.5), Wilcoxon rank-sum test). In samples from mice infected with *A. fumigatus* Δ*pptA*, 24 metabolites including methionine sulfoxide showed an increased abundance and 9 metabolites showed a decreased abundance (Figure 3B, Table S3). As methionine sulfoxide is formed by oxidation of the sulfur atom of methionine by reactive oxygen species (ROS) and therefore serves as an antioxidant ^24^, an increase in its abundance could be linked to ROS production during response of the residual immune system in immune suppressed mice to the infection of the lungs.

As we observed changes due to *A. fumigatus* infection in all four analyzed sample types, *i.e*., the gut microbiome, lung microbiome, plasma metabolome and lung metabolome, we further analyzed the communication axis between gut and lung. We calculated the global mediation effect for significantly different abundant metabolites and genera (*p* < 0.05, |log_2_FC| > log_2_(1.5), Wilcoxon rank-sum test for metabolites, DESeq2 for genera, the union of CA *vs* WT and CA *vs* pptA) between lung and gut through the plasma metabolome. For infection with *A. fumigatus*, there was a statistically significant global influence of the gut microbiome on the lung metabolome through the plasma metabolome (*p* = 0.0011 for *A. fumigatus* wild-type infection; *p* = 0.0014 for *A. fumigatus* Δ*pptA* infection, Wilcoxon rank-sum test). Only an infection with *A. fumigatus* wild type showed a significant global mediation effect of lung metabolome on the gut microbiome (p < 0.05, Wilcoxon rank-sum test) (Figure 4C). Although there are no global mediation effects between gut microbiome and lung microbiome through the plasma metabolome, 3-hydroxybutyric acid and crotonic acid were found to mediate specific interactions between gut and lung bacteria. *Ligilactobacillus*, the most abundant genus in the lung microbiome, showed a significant mediation effect on the gut bacterium *Dysosmobacter* by crotonic acid, whereas the gut bacterium *Limosilactobacillus* had a significant effect on the lung bacterium *Ligilactobacillus* mediated by 3-hydroxybutyric acid. In summary, we found that infection with wild-type *A. fumigatus* led to dysbiosis of the gut microbiome, which potentially induced changes in the plasma and lung metabolomes. Although the infection with wild-type *A. fumigatus* had a smaller effect on the lung microbiome, it caused significant increase of relative abundance of the most abundant genus *Ligilactobacillus.* Infection with the avirulent Δ*pptA* strain had comparatively minor effects on the microbiomes but equally pronounced effects on the metabolomes as an infection with the wild-type of *A. fumigatus*.

**Figure 4:**
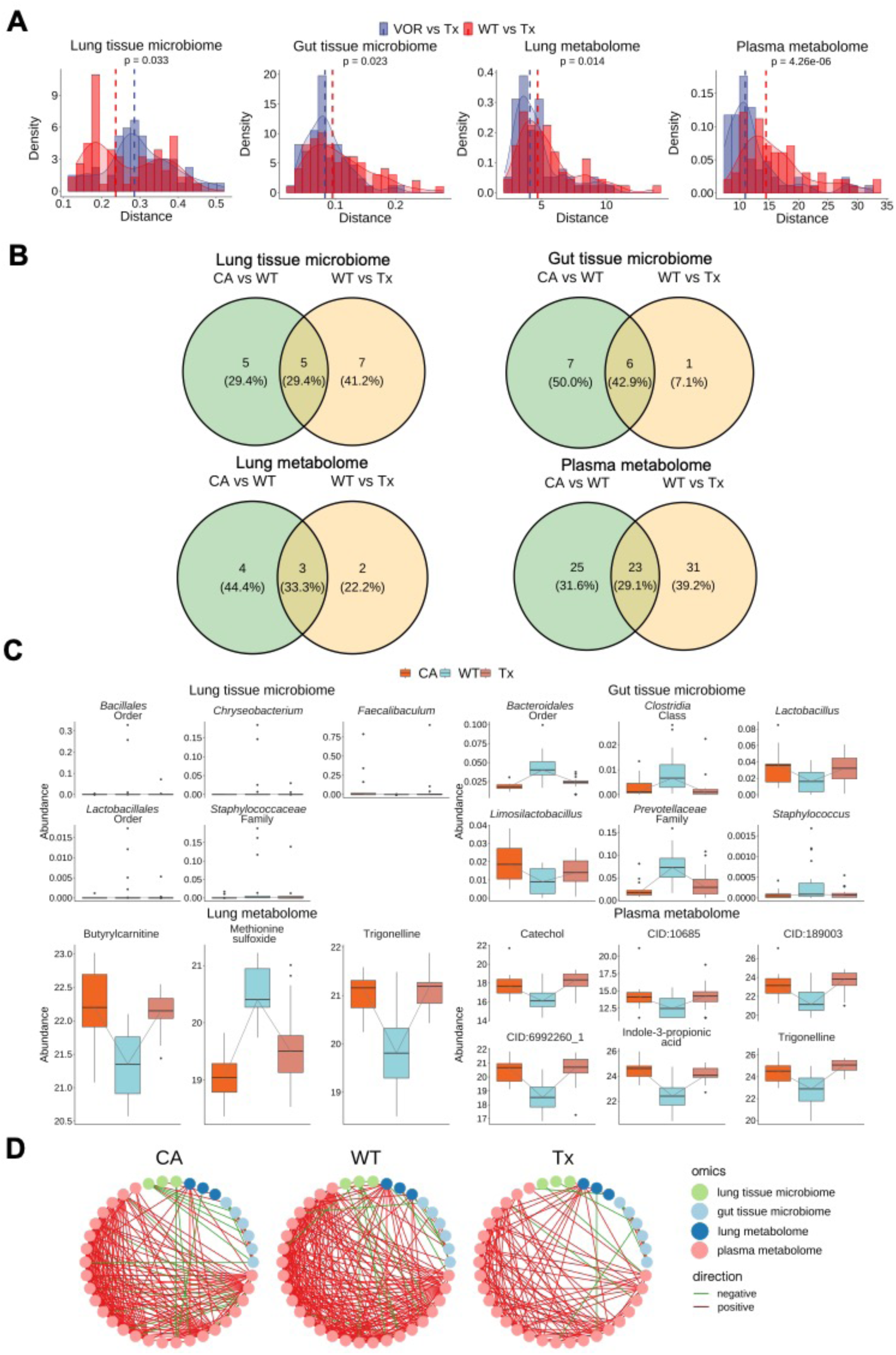
Voriconazole treatment after infection with *A. fumigatus* reduces alterations of the lung microbiome, gut microbiome, lung metabolome and plasma metabolome. Immunosuppressed mice infected with *A. fumigatus* and treated with voriconazole (Tx) were compared with immunosuppressed mice infected with *A. fumigatus* (WT) and with immunosuppressed mice treated with voriconazole (VOR). A) Sample distances of WT and VOR to Tx, respectively (weighted UniFrac for microbiome and Euclidean for metabolome, *p*-values of density differences calculated by Wilcoxon test). B) Venn diagrams of differentially abundant genera and metabolites (*p* < 0.05, |log_2_FC| > log_2_(1.5)) for *A. fumigatus* infection (CA *vs*. WT) and voriconazole treatment (CA *vs*. Tx). C) Abundance of differentially abundant genera and metabolites that exhibit a shift in abundance direction for CA *vs*. WT and WT *vs*. Tx. D) Significant correlations (*p* < 0.05, Spearman’s correlation) between genera and metabolites with differential abundance and partial recovery after antifungal treatment. For Figures B-D bacteria and metabolites directly influenced by voriconazole treatment were not considered in the analysis. Abbreviations: CID:10685: 3-(2-Hydroxyethyl)indole; CID:189003: 3,4-Dihydroxybenzenesulfonic acid; CID:6992260: 3-(Propan-2-yl)-octahydropyrrolo[1,2-a]pyrazine-1,4-dione.

### Antifungal treatment partially inhibits alterations of microbiomes and metabolomes

Next, we tested whether antifungal treatment with voriconazole can reduce major changes of the microbiome when given at the onset of infection with *A. fumigatus*. Therefore, the inter-group beta diversity of the lung and gut microbiome (weighted UniFrac distance) and the lung and plasma metabolome (Euclidean distance) were compared between infected voriconazole-treated mice and untreated mice infected with wild-type *A. fumigatus*, or mice that were not infected but were only treated with voriconazole. The gut microbiome, lung metabolome and plasma metabolome of infected and voriconazole-treated mice were found to be significantly closer to uninfected voriconazole-treated mice (*p* < 0.05, Wilcoxon rank-sum test), while the lung microbiome of infected voriconazole-treated mice was significantly closer to infected mice (Figure 4A), but the presence of two peaks suggests that the lung microbiome may require a longer recovery period (Figure 4A).

We evaluated the significantly changed bacteria and metabolites after wild-type *A. fumigatus* infection and antifungal treatment and observed that the voriconazole treatment also reduced the effect of *A. fumigatus* wild type infection on the levels of individual bacteria and metabolites (Figure 4B, 4C, Tables S1-S4). In the lung microbiome, the change in relative abundances of 5 genera caused by wild-type *A. fumigatus* infection did not occur during voriconazole treatment. Although the voriconazole treatment did not completely prevent changes in relative abundance of *Ligilactobacillus*, its abundance was lower compared to infected untreated mice (*p* = 0.674, FDR = 0.701, log_2_FC = -0.420, DESeq2). The same effect of the voriconazole treatment was observed for 6 genera in the gut, including abolished decrease of *Lactobacillus*, and *Limosilactobacillus*, and abolished increase of *Staphylococcus.* Notably, relative abundance of *Escherichia*/*Shigella* was not statistically significantly reduced (*p* = 0.09) in the gut of infected and voriconazole treated mice compared to infected and not treated mice. For the metabolomes, change for 3 lung metabolites and 6 plasma metabolites was at least partially prevented by voriconazole treatment of infected mice. Interestingly, methionine sulfoxide was statistically significantly reduced in the lung of infected and voriconazole treated mice compared to infected untreated mice, suggesting less ROS production in the lungs of infected voriconazole treated mice ^24^.

Furthermore, a correlation network that was created to investigate interactions between the differentially abundant genera and metabolites showed that the interactions increased after wild-type *A. fumigatus* infection but not after *A. fumigatus* infection and voriconazole treatment (Figure 4D). These findings are consistent with the mediation analysis results, which indicated an increased mediating effect of the gut microbiome on the lung metabolome through the plasma metabolome (Figure 3C). Therefore, these interactions were not observed in samples where *A. fumigatus* was eliminated by antifungal treatment.

### *Ligilactobacillus murinus* abundance increases in lungs infected with *A. fumigatus*

Since we observed a statistically significant increase in the relative abundance of *Ligilactobacillus* in lungs infected with *A. fumigatus* and this bacterium was most frequently detected in all samples, we quantified its absolute abundance in the lung samples by quantitative PCR and tested whether it correlates with the *A. fumigatus* burden. DNA of *L. murinus* could be detected in 3 out of 14 samples of immunosuppressed mice, 10 out of 20 samples of mice infected with *A. fumigatus* wild type, 5 from 18 samples of mice infected with *A. fumigatus* wild type and treated with voriconazole, and 4 out of 20 samples of mice infected with the *ΔpptA* strain. In samples in which we did not detect *L. murinus*, its DNA content was apparently below the detection limit and/or the preparation might have failed due to the cell wall of the gram-positive bacterium.

The absolute abundance of DNA of *L. murinus* was increased in the lung samples infected with *A. fumigatus* wild type compared to both, uninfected immunocompromised mice (*p* = 0.075, Wilcoxon rank-sum test) and immunocompromised mice infected with *A. fumigatus* wild type and treated with voriconazole (*p* = 0.057, Wilcoxon rank-sum test) (Figure 5A). Moreover, when the samples with *L. murinus* detected by qPCR were analyzed, a positive correlation (*R* = 0.725, *p* = 0.003, Spearman’s correlation) between *L. murinus* and *A. fumigatus* burden in infected lungs was observed (Figure 5B).

**Figure 5:**
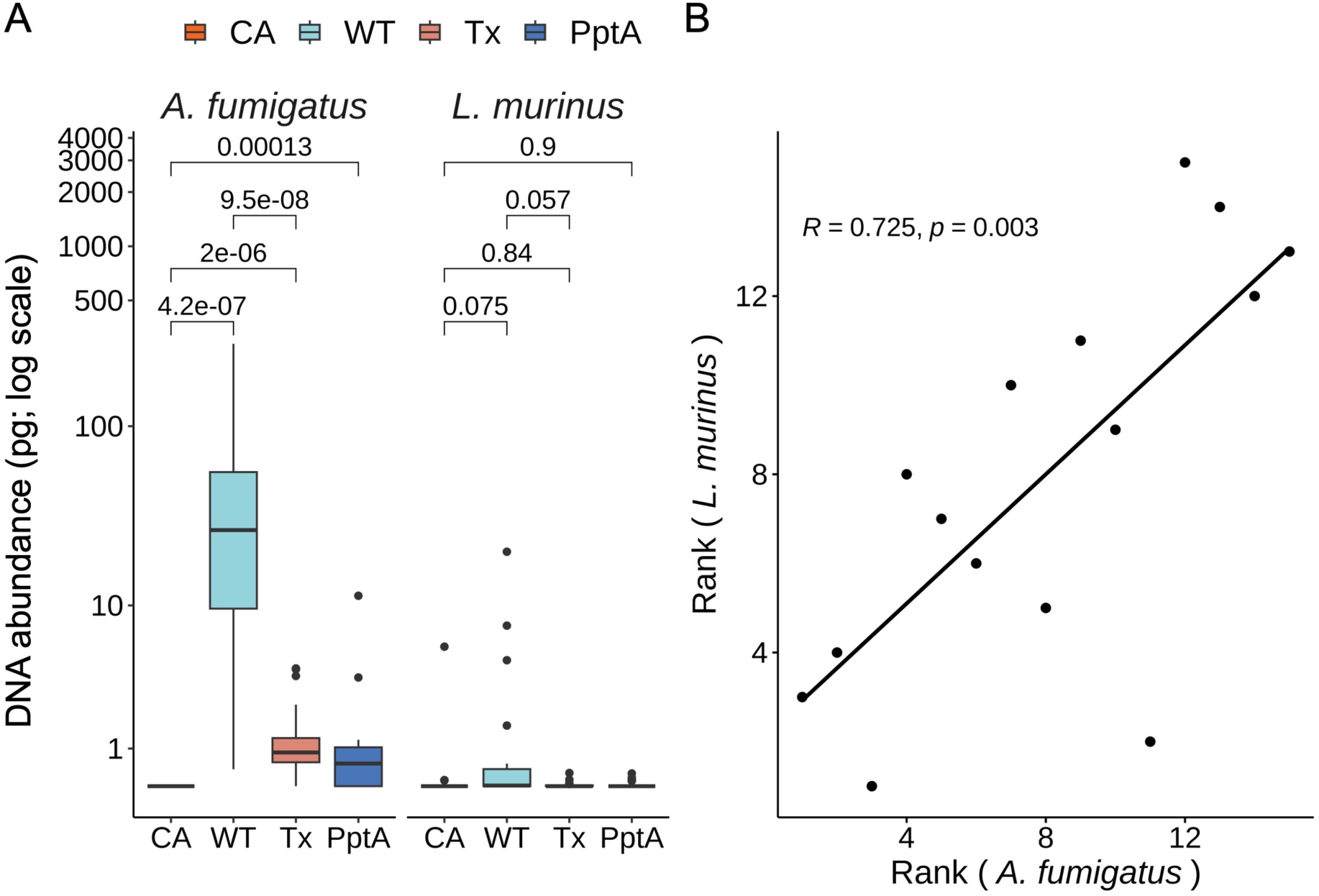
*L. murinus* increases in abundance in lungs infected with *A. fumigatus* and this correlates with the fungal burden. A) *L. murinus* and *A. fumigatus* quantification by qPCR. DNA quantity in pg per 1 µg of mouse DNA is shown. Wilcoxon rank-sum test was used to calculate the *p*-values. B) Correlation of *L. murinus* and *A. fumigatus* burden in infected lungs. The Spearman’s correlation was used to determine the abundance association between *A. fumigatus* DNA and *L. murinus* DNA, only samples with detected *L. murinus* DNA were included in the analysis.

### *L. murinus* is among the most prominent bacteria cultivated from the lower respiratory tract of mice

Only a few studies have been performed to detect live bacteria in both the human and mouse lung ^25–30^ and in particular to relate DNA sequencing data with culturable bacteria from the lung of mice. To validate lung microbiome results analyzed by 16S rDNA sequencing, we isolated viable bacteria from lung and tracheal tissue samples of C57BL/6 mice. Bacteria were isolated from 71.4 % of the lung samples and from all tracheal samples. The bacterial diversity was very similar between the tracheal and lung samples, however, more colony forming units (CFUs) were observed in the tracheal samples than in the lung samples (Figure S3). The most frequently isolated species, from both sites, include *Ligilactobacillus murinus*, *Lactobacillus taiwanensis*, *Streptococcus danieliae, Staphylococcus xylosus*, *Limosilactobacillus reuteri*, and *Lactobacillus intestinalis* (Figure 6A). The majority of these species also belonged to the most abundant genera in lung microbiome data obtained by 16S rDNA sequencing (Figure S1A).

**Figure 6:**
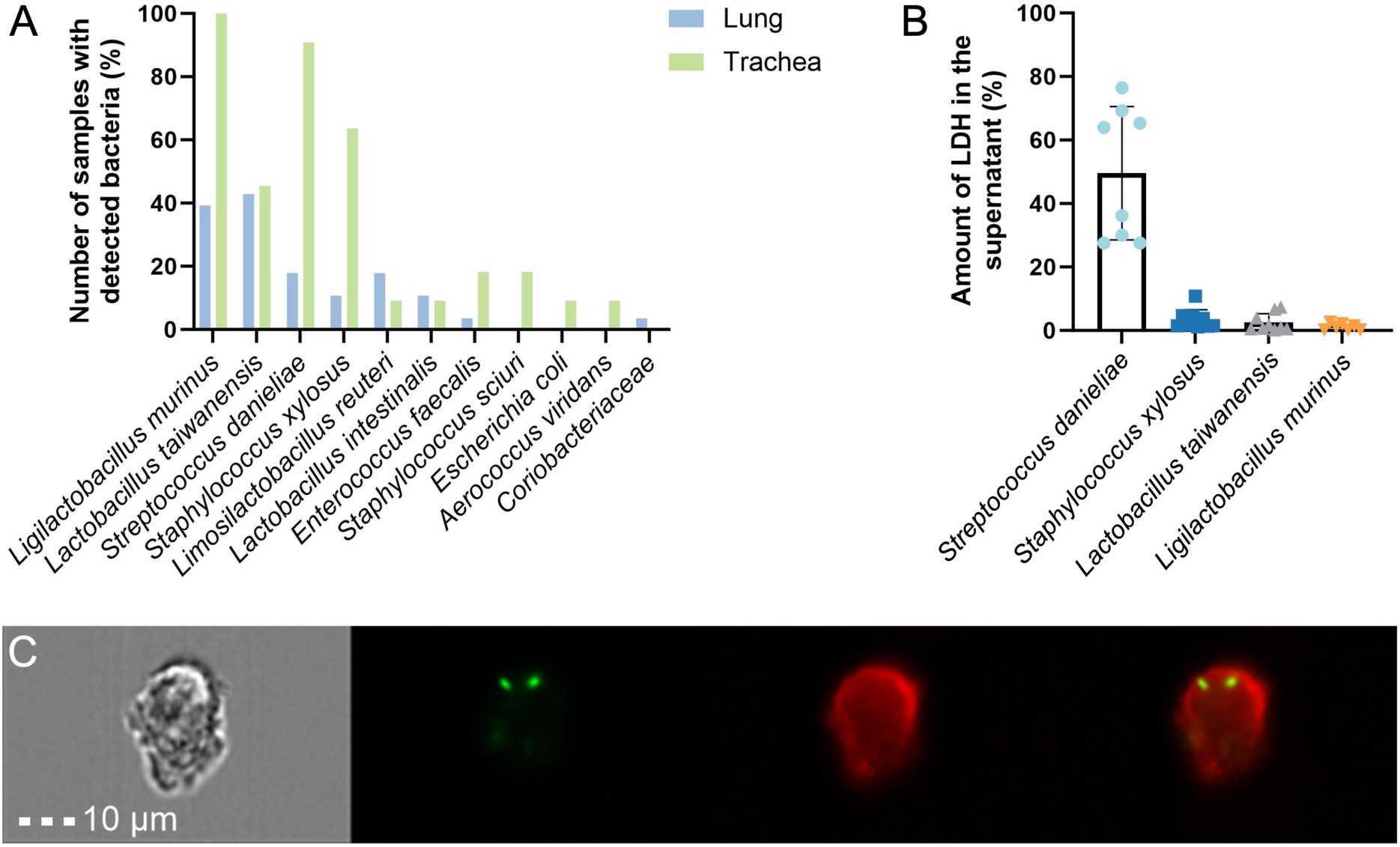
Culturable bacteria isolated from the mouse lower respiratory tract show different interactions with alveolar epithelial cells. A) Bacteria isolated from lung and trachea samples; percentage of samples positive for each bacterium is shown. Twenty-eight lung and 11 tracheal samples were used for bacteria isolation. B) Cytotoxicity of *S. danieliae*, *S. xylosus*, *L. taiwanensis* and *L. murinus* on mouse T7 alveolar epithelial cells determined by quantification of released lactate dehydrogenase (LDH) after 24 hours of infection. 0 % cytotoxicity: spontaneous LDH release (non-treated cells); 100 % cytotoxicity: maximal LDH release (cells treated with lysis buffer). Error bars represent the standard deviation. C) Analysis of internalization of GFP-producing *L. murinus* (green) by mouse alveolar epithelial cells stained with concanavalin-AF647 (red) six hours after infection. The analysis was performed using imaging flow cytometry.

### *L. murinus* from the lower respiratory tract of mice is tolerated and even internalized by alveolar epithelial cells

Alveolar epithelial cells serve as the first barrier in the alveoli, protecting the host from microbial invasion ^31^. Because of their presence in the lung of mice, *L. taiwanensis*, *L. murinus*, *S. danieliae* and *S. xylosus* were analyzed for their interaction with these host cells. Therefore, we assessed whether these bacteria cause cell damage of murine alveolar epithelial cells or are tolerated by the cells. Whereas infection with *S. danieliae* resulted in increased cell damage, *L. murinus*, *L. taiwanensis* and *S. xylosus* caused no obvious damage to epithelial cells (Figure 6B).

We further investigated whether lung bacteria can be internalized by alveolar epithelial cells which can also phagocytose particles and pathogens, although they are not professional phagocytes ^32^. We focused on the internalization of *L. murinus* because (i) it was one of the most abundant bacteria detected both by sequencing and isolation approaches in our experiments and also in previous studies ^33,34^ and furthermore, (ii) *L. murinus* did not damage epithelial cells. To visualize the internalization of *L. murinus* by epithelial cells, we generated GFP-producing bacteria. Six hours after infection, 1.2 ± 0.17 % of mouse T7 alveolar epithelial cells contained internalized GFP-producing *L. murinus* (Figure 6C) which is in a similar range as the uptake of *A. fumigatus* conidia ^32^. This shows that *L. murinus* isolated from the murine lower respiratory tract can be internalized at a low rate by murine alveolar epithelial cells.

### *A. fumigatus* creates a microaerophilic habitat that promotes the growth of *L. murinus*

Since our data suggest that infection with *A. fumigatus* leads to an increase in the abundance of in particular anaerobic bacteria, most prominently of *L. murinus*, we investigated *in vitro* whether there is a direct effect of the fungus on the bacteria. Therefore, we co-cultured *A. fumigatus* with the *L. murinus* strain MAM1902 isolated from the mouse lower respiratory tract and observed that the number of bacterial CFUs was significantly higher in co-cultures (Figure 7A, Figure S4A). Further, bacterial growth occurred in close proximity to the fungal hyphae (Figure 7B, 7C).

**Figure 7:**
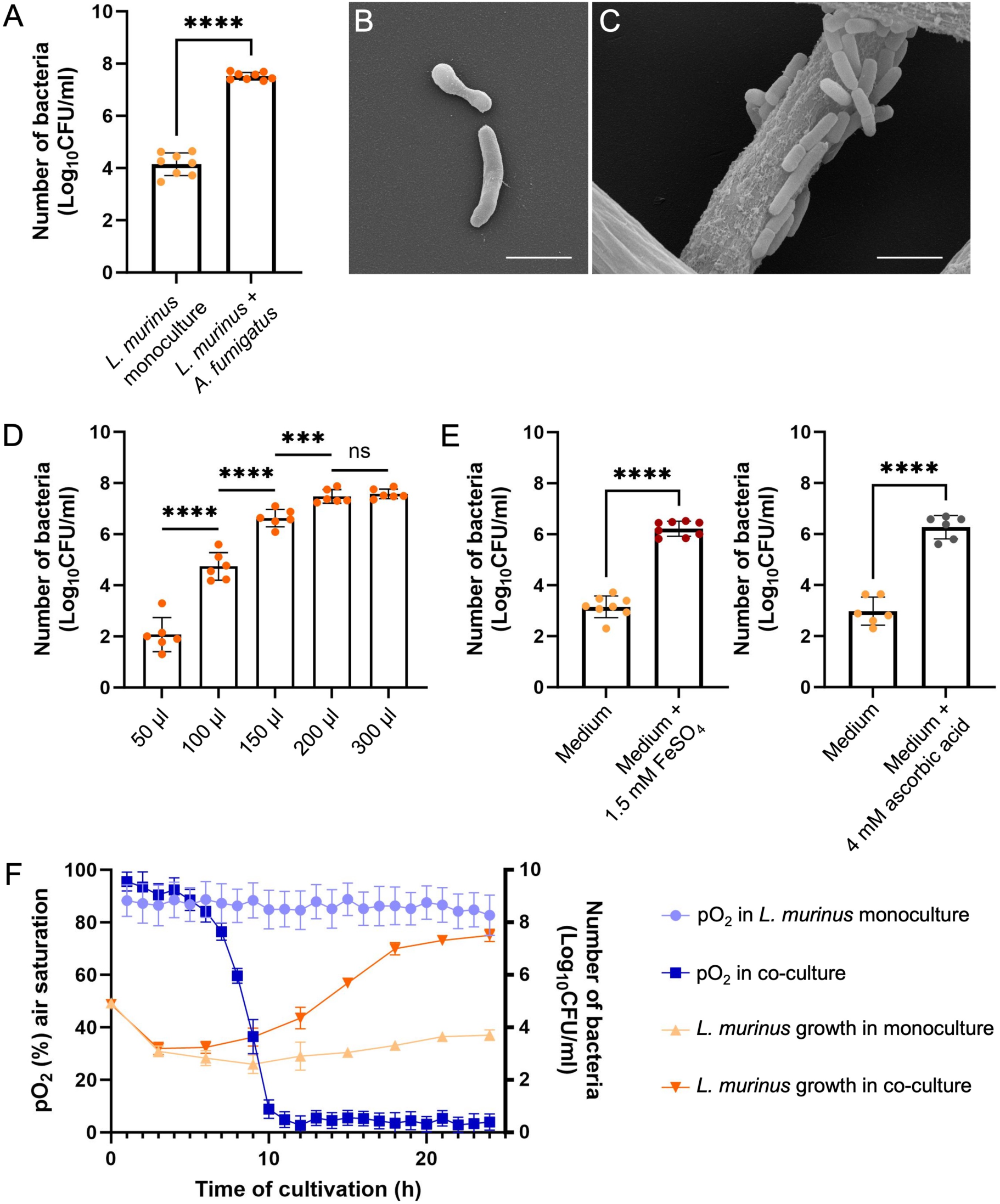
*A. fumigatus* promotes the growth of *L. murinus* by creating a microaerophilic environment. A) Number of CFUs of *L. murinus* monoculture and co-culture with *A. fumigatus* after 24 hours of cultivation. B,C) Scanning electron microscopy of *L. murinus* in monoculture (B) and co-culture with *A. fumigatus* (C) after 24 hours of cultivation; scale bar, 2 µm. D) Dependence of *L. murinus* growth in co-culture with *A. fumigatus* on culture volume; cultivation was carried out in 96-well plates for 24 h. ns, not significant. E) CFUs determined from *L. murinus* cultured with and without supplementation of ferrous sulfate and ascorbic acid. F) Correlation between pO_2_ and *L. murinus* growth in monoculture and co-culture with *A. fumigatus* over 24 h. Cultivation was performed in Ham’s F-12K medium. Unpaired t test was used to assess the statistical significance (****, *p* < 0.0001; ***, *p* < 0.001; **, *p* < 0.01; *, *p* < 0.05).

Next, we investigated the mechanistic basis of this growth-promoting effect. By co-culturing *L. murinus* with various *A. fumigatus* mutant strains we observed that growth promotion by *A. fumigatus* was independent of TAFC siderophore production, the production of other natural products or the reduction of Fe^3+^ to Fe^2+^ by the surface iron reductase FreB (Figure S4B). Another hypothesis we tested was the consumption of oxygen by the fungus and thus the creation of a hypoxic environment along the hyphae that could promote growth of the anaerobic bacteria. When using different media volumes (50 µl to 300 µl) for co-cultivation in 96-well plates, we observed that the growth-promoting effect depended on the culture volume, *i.e*., the number of CFUs increased with increasing the volume from 50 µl to 200 µl, but did not increase further from 200 µl to 300 µl (Figure 7D). One reason could be that the lower surface to volume ratio leading to reduced gas exchange in higher volumes resulted in a hypoxic environment promoting the growth of the anaerobic bacteria. By measurement of the oxygen concentration, we confirmed oxygen depletion of the medium in the presence of *A. fumigatus*, leading to anoxia within 10 hours (Figure 7F, Figure S4C). By contrast, oxygen saturation of *L. murinus* mono-cultures remained at around 80-100 % over 24 hours. The reduction of oxygen concentration correlated with increased growth of *L. murinus* in co-culture (Figure 7F). Addition of the oxygen scavengers ascorbic acid and ferrous sulfate to the medium also enhanced *L. murinus* growth (Figure 7E). Collectively, oxygen consumption by *A. fumigatus* creates a hypoxic environment that promotes the growth of anaerobic *L. murinus* in co-cultures.

## Discussion

Our results indicate that lung infection of mice with *A. fumigatus* not only directly influences the lung microbiome and metabolome but also the gut microbiome, which is apparently triggered by the change of the lung and plasma metabolome. The alteration in the gut microbiome was most prominent for 13 genera showing a change in their abundances, most notably *Escherichia/Shigella*. *Escherichia/Shigella* are considered to be facultative pathogens and frequently increase in abundance of these bacteria is a sign of dysbiosis and indicates reduced immune protection ^22,23^ that is caused here by immune suppression coupled with infection. *Ligilactobacillus* was the most abundant genus in the lung microbiome and might be considered as a member of the lung microbiome. Based on our correlation analysis, *Ligilactobacillus* showed a significant mediation effect on the gut bacterium *Dysosmobacter* by crotonic acid, whereas the gut bacterium *Limosilactobacillus* had a significant effect on the lung bacterium *Ligilactobacillus* mediated by 3-hydroxybutyric acid. 3-hydroxybutyric acid and crotonic acid are short chain fatty acids that are intermediates of the fatty acid biosynthesis that occurs in both bacteria and mammals ^35^. Short chain fatty acids have been shown to be key mediators affecting the host in various ways ^36^. Since supplementation with 3-hydroxybutyric acid did not influence the growth of *L. murinus in vitro* (data not shown), the observed mediation effect was most probably indirect. Collectively, the findings that *A. fumigatus* infection leads to gut dysbiosis are surprising given that *A. fumigatus* is infecting *via* the lung. These findings might have consequences for future therapeutic strategies considering the established role of the gut microbiome in regulating immune responses. Furthermore, the correlation of changes in the gut microbiome and plasma metabolome and the observed influence on changes in the lung metabolome by the gut microbiome *via* the plasma metabolome indicate the importance of gut bacteria for lung homeostasis and support previous findings ^37–39^.

Immunosuppression is crucial for the study of invasive aspergillosis, as the immune system of an immunocompetent host usually prevents systemic fungal infection. Various patient groups including those receiving high-dose corticosteroids, however, are at a high risk for this fungal infection ^13^. Previous studies on mice have shown that corticosteroid treatment affects the gut microbiome ^40–42^. Our results are consistent with these studies because we showed that immunosuppression led to dysbalance of the gut microbiome and drastic changes in the plasma and lung metabolome, whereas for the lung microbiome, we did not observe significant differences in the alpha and beta diversity. However, abundance of 8 genera significantly differed, with the largest change seen for *Bifidobacterium,* but also others like *Faecalibaculum*, *Staphylococcaceae* family, *Anaerococcus*, *Methanothermobacter*, *Lactobacillus*, *Photobacterium*, and *Latilactobacillus*. These bacteria might be controlled by a healthy immune system either as constituents of the lung microbiome or when they were inhaled. This conclusion is underlined by previous results demonstrating that the abundance of *Staphylococcus*, *Lactobacillus* and *Anaerococcus* was shown to be increased during lung disorders ^1^, in the lungs of patients with reduced lung function ^43^, and in the upper respiratory tract of patients with chronic rhinosinusitis, respectively ^44^.

After an infection of mice with the avirulent *A. fumigatus ΔpptA* mutant, the lung microbiome was much less affected than after infection with the virulent wild-type strain. Similarly, treatment of *A. fumigatus* wild-type infection with voriconazole also led to changes of the microbiome, however, these changes were much less pronounced and similar to those seen for infections with the *ΔpptA* mutant strain. The reason might be that in both cases conidia of *A. fumigatus* could not germinate and produce mycelia in the lung, as shown before for the *ΔpptA* mutant strain ^21^, and thus do not lead to a dysbiosis as seen for the infection with the wild-type strain. Since we administered the antifungal drug to the mice immediately after the onset of the infection, this could have prevented the germination of the conidia and formation of hyphae and thus the establishment of the infection. This conclusion on voriconazole is also supported by the analysis of the gut microbiome, plasma metabolome and lung metabolome that were close to a state of uninfected mice when infected mice were treated with voriconazole. At least the data generated here in the mouse infection model, supports the value of a preemptive therapy ^45,46^, given the antifungal drug is administered early. The effect of voriconazole treatment of uninfected mice was not as pronounced as in previous studies ^47,48^, which might be due to the lower dosage and shorter treatment duration applied here, as well as the imbalance caused by immunosuppression.

There is evidence from *in vitro* experiments and *in vivo* analyses that *A. fumigatus* can modulate the microbiota ^49–51^. For example, in the lung of cystic fibrosis patients, *A. fumigatus* apparently shapes a lung microbiome towards a more beneficial environment for fungal growth that is associated with aromatic amino acid availability and the shikimate pathway ^50^. It is conceivable that for shaping the microbiome the fungus also uses its defensin-like peptides on the conidial surface and its natural products that exhibit antimicrobial and toxic activity ^49,51,52^. In the lung of mice, it was shown that *A. fumigatus* can produce several natural products including the toxin gliotoxin ^53^. Although not analyzed yet in a mammalian microbiome, *A. fumigatus* has the ability to produce several other natural products like fumicyclines and fumigermins that have antibacterial activity ^54,55^. In line with these considerations, we observed a decrease in relative abundance for some of the bacteria detected in the infected lungs that could be due to factors like natural products, defensins and/or the residual activity of the immune system.

It was striking to observe that infection of immunosuppressed mice with *A. fumigatus* wild type led to a higher abundance of seven genera, two of them are composed of obligate anaerobic species and three genera of facultative anaerobic species, most prominently of *Ligilactobacillus*, which was not observed in lung samples infected with the avirulent *ΔpptA* strain. *L. murinus* was also previously detected in the mouse lower respiratory tract by both sequencing and isolation ^33,34,56^. In further *in vitro* experiments, we demonstrated that there is a direct increase of the CFUs of *L. murinus* in presence of *A. fumigatus*. The most plausible explanation for this interesting direct interaction, was the creation of a microaerophilic environment through oxygen consumption by the fungus. This hypothesis was underlined by our findings that *L. murinus* grew much better under anaerobic conditions and that during cultivation of *A. fumigatus* in mono-culture and co-cultivation with *L. murinus*, the oxygen concentration in the culture medium decreased overtime to 0 %. Moreover, it is known that oxygen consumption by aerobic microorganisms can facilitate or prevent colonization by certain microorganisms both in soil and in the host. For example, the commensal fungus *Candida albicans* can promote the growth of anaerobic bacteria in co-cultures through the consumption of dissolved oxygen ^57,58^. Similarly, the fungus *Coprinopsis cinerea*, which lives in oxic soil habitats, promotes the germination and growth of obligate anaerobic *Clostridium acetobutylicum* bacteria ^59^. In a recent clinical study, an increase in strict anaerobes belonging to the *Clostridiales* was observed in bronchoalveolar lavage samples from immunocompromised patients with invasive aspergillosis ^10^. It is thus conceivable that *A. fumigatus* promotes the growth of microaerophilic and likely anaerobic bacteria even in the lung during infection by creating hypoxic niches when the fungus germinates and grows by both the consumption of oxygen but also by triggering inflammation. The latter was also shown to lead to hypoxia, for example during acute lung injury in the lung, or bowel disease in the intestine ^60–62^. All these examples support the hypothesis that the availability of oxygen is an important factor for shaping a microbiome. It can be manipulated by aerobic microorganisms that consume oxygen and lead to necrosis of tissue.

In summary, our results suggest that *A. fumigatus* shapes its lung microbiome by creating a microaerophilic or even anaerobic environment during infection thereby promoting the growth of anaerobic bacteria most prominently of *L. murinus,* which is intuitively contradictory what would be expected in the lung environment. This finding could lead to a potential use of detection of certain anaerobic bacteria in the lung microbiome for the diagnosis of *A. fumigatus* infection. However, it remains to be shown, whether or how these bacteria influence the infection. In *Mycobacterium tuberculosis*-infected mice, administration of a specific *L. murinus* strain resulted in reduction of pulmonary inflammation, without increasing *M. tuberculosis* burden. It was suggested that this strain may shape local T cells in mice and may play a protective role against *M. tuberculosis* ^34^. *L. murinus* can even directly affect pathogens as shown for *S. pneumonia* whose growth was inhibited *in vitro* through the production of lactic acid ^33^. At least, we showed that *L. murinus* is apparently not toxic to lung epithelial cells, supporting its presence in the lung. Furthermore, by generating a GFP-producing *L. murinus* we observed internalization of bacteria in lung epithelial cells in a low rate which might influence the immune response against pathogens.

## Supporting information

Supplementary Table S5 Weight & Scoring

Supplementary Table Metabolites Area under the curve Lung

Supplementary Table Metabolites Area under the curve Plasma

## Acknowledgments

We thank Sigrun Kirste, Andrea Hartmann, Sylke Fricke, and Christina Täumer for excellent technical assistance. We thank Dr. Olaf Kniemeyer for the help in the project administration. This work was funded by the Deutsche Forschungsgemeinschaft (DFG) cluster of excellence *Balance of the Microverse* (Project-ID 390713860, Gepris 2051), DFG CRC/Transregio 124 ‘FungiNet (projects A1, C5, INF; 210879364), and EU-funded Horizon 2020 project HDM-FUN (ID 847507). L.N. was also supported by the DFG excellence graduate school *Jena School for Microbial Communication*. G.P was also supported by the German Federal Ministry of Education and Research within the funding project PerMiCCion (project ID: 01KD2101A).

## Author Contributions

Conceptualization, A.A.B. and G.P.; methodology, L.N., X.C., L.R., M.S., S.N., T.H., I.J., G.P., and A.A.B.; formal analysis, L.N., X.C., I.J., G.P., and A.A.B.; investigation, L.N., X.C., L.R., K.L., B.S., S.N., and T.H.; resources, M.S., W.B., S.V., T.H., I.J.; writing – original draft, L.N., X.C., T.H., and A.A.B.; writing – review & editing, L.N., X.C., B.S., M.S., S.N., T.H., I.J., G.P. and A.A.B.; visualization, L.N., X.C.; supervision, G.P. and A.A.B.; project administration, L.R., G.P., and A.A.B.; funding acquisition, A.A.B. and G.P..

## STAR Methods

## RESOURCE AVAILABILITY

### Lead Contact

Further information and requests for resources and reagents should be directed to and will be fulfilled by the lead contact, Axel A. Brakhage.

### Materials Availability

- All materials within the paper are available from the corresponding author upon reasonable request.
- This study did not generate new unique reagents.

### Data and Code Availability

- The metagenome data have been deposited at the European Nucleotide Archive (project ID: PRJEB76136).
- Isolated strains were deposited at the Jena Microbial Resource Collection (https://www.leibniz-hki.de/de/jena-microbial-resource-collection.html)
- The source code and necessary data for results generated in this manuscript are avail-able at github: https://github.com/codechenx/release_mice_aspergillus_infec-tion_lung_project.
- Any additional information required to reanalyze the data reported in this work paper is available from the Lead Contact upon request.

## EXPERIMENTAL MODEL AND SUBJECT DETAILS

### Mouse model of invasive aspergillosis

To induce invasive aspergillosis in mice, we applied an established infection model using BALB/cByJRj mice. In inbred mice, the inter-individual variation in the core microbiome is much lower compared to outbred mice ^63^. Female mice (8 weeks) were purchased from Janvier Labs and housed under standard SPF conditions. The acclimatization phase of the animals was 1 week. 105 mice in six treatment groups were included in the experiment: PBS-control mice, PBS-control mice immunosuppressed with cortisone acetate, PBS-control mice immunosuppressed with cortisone acetate and treated with voriconazole, immunosuppressed mice infected with *A. fumigatus* wild type, immunosuppressed mice infected with *A. fumigatus* Δ*pptA* mutant, and immunosuppressed mice infected with *A. fumigatus* wild type and treated with voriconazole (Figure 1A). For immunosuppression, mice were treated intraperitoneally with cortisone acetate (1 g/kg) on the days -3 and 0 of the experiment. Infection with 2 x 10^5^ *A. fumigatus* A1160^+^ wild type or Δ*pptA* conidia was performed intranasally on day 0. Voriconazole treatment (40 mg/kg) was performed intravenously on day 0 (directly after infection), day 1, and day 2. On day 3, mice were euthanized, cecum and lung samples were obtained under sterile conditions and kept at -70 °C until further processing. Blood from euthanized mice was drawn directly from the heart *via* puncture and then centrifuged at 2000 *g* for 15 min. Fifty microliters of blood plasma were stored at -70 °C until extraction. In addition to the samples from the cecum, we also collected feces on day -5 to -4 (before immunosuppression) and day -1 (after immunosuppression). Here, the animals were placed in a separate cage for a maximum of 90 minutes. The 3-4 fecal grains were then collected.

All animals were handled in compliance with the European animal welfare regulations. Approval for the experiments was obtained from the responsible federal/state authority and the ethics committee, which are based on the provisions of the German Animal Welfare Act (permit no HKI-21-008).

## METHOD DETAILS

### Lung sample preparation

To obtain uniform samples for subsequent metagenomic and metabolomic analyses, frozen lung samples were homogenized in ice-cold DNA-free water (Qiagen) to the final concentration of 66.7 mg/ml (w/v) using a gentleMACS Dissociator (Miltenyi Biotec). Homogenized samples containing 50 mg tissue were directly frozen at -70 °C until extraction of DNA and metabolites.

### DNA extraction and sequencing from lung samples

Sample analysis was carried out by DNASense (Aalborg, Denmark) as follows: 375 μl of mouse lung homogenate (∼25 mg tissue) from each sample were used for DNA extraction using the Ultra-Deep Microbiome kit according to the manufacturer’s recommendations (Molzym) with modifications: initial proteinase K digestion was extended to 20 min, centrifugation steps were at 15,000 *g*, and final elution was done with 40 µL deionized water heated to 70 °C. After depletion of the host DNA, the pellet was resuspended in 1 ml RS buffer and half of the volume was used for PCR amplification and sequencing of the bacteria/archaea 16S rRNA gene variable region 4 (abV4-C). Gel electrophoresis using Tapestation 2200 (Genomic DNA and D1000 screentapes, Agilent) was used to validate product size and purity of a subset of DNA extracts. DNA concentration was measured using a Qubit dsDNA HS Assay kit (Thermo Fisher Scientific).

Amplicon libraries for the bacteria/archaea 16S rRNA gene variable region 4 (abV4-C) were prepared according to an Illumina protocol. Up to 10 ng of extracted DNA was used as template for PCR amplification of the taxonomic marker genes for 30 cycles of amplification. Duplicate PCR reactions were performed for each sample and the duplicates were pooled. The forward- and reverse-tailed primers targeting the bacteria/archaea 16S rRNA gene variable region 4 (abV4-C) were 515FB (GTGYCAGCMGCCGCGGTAA) and 806RB (GGACTACNVGGGTWTCTAAT), respectively ^64,65^. The primer tails enable annealing of Illumina Nextera adaptors required for sequencing in a subsequent PCR. The resulting amplicon libraries were purified using the standard protocol for CleanNGS SPRI beads (CleanNA) with a bead to sample ratio of 4:5. DNA was eluted in 25 μL of nuclease-free water (Qiagen). Sequencing libraries were prepared from the purified amplicon libraries using a second PCR for 8 cycles. The resulting sequencing libraries were purified using the standard protocol for CleanNGS SPRI beads with a bead to sample ratio of 4:5. DNA was eluted in 25 μL nuclease-free water. The purified sequencing libraries were pooled in equimolar concentrations and diluted to 2 nM. The samples were paired-end sequenced (2 x 300 bp) on a MiSeq (Illumina) using a MiSeq Reagent kit v3 (Illumina) following the standard guidelines for preparing and loading samples on the MiSeq. PhiX control library consisting of balanced base composition (∼45 % GC and ∼55 % AT) was spiked in at > 10 % for the sequencing control (DNASense, Arlborg, Denmark).

Amplicon libraries for the bacteria 16S rRNA gene variable regions 1-8 (bV18-A) were prepared using a custom protocol. Up to 25 ng of extracted DNA was used as template for PCR amplification, and each PCR reaction (50 μL) contained 0.5 mM dNTP mix, 0.01 units of Platinum SuperFi DNA Polymerase (Thermo Fisher Scientific, USA), and 500 nM of each forward and reverse primer in the supplied SuperFI Buffer. PCR was done with the following program: Initial denaturation at 98 °C for 3 min, 28 cycles of amplification (98 °C for 30 s, 62 °C for 20 s, 72 °C for 2 min) and a final elongation at 72 °C for 5 min. The forward and reverse primers used included custom 24 nucleotide barcode sequences followed by the sequences targeting bV18-A and were follows: 8F (AGRGTTYGATYMTGGCTCAG) and 1391R (GACGGGCGGTGWGTRCA) ^66^. The resulting amplicon libraries were purified using the standard protocol for CleanNGS SPRI beads (CleanNA, Netherlands) with a bead to sample ratio of 3:5. DNA was eluted in 25 μL of nuclease free water (Qiagen, Germany). Sequencing libraries were prepared from the purified amplicon libraries using the SQK LSK114 kit (Oxford Nanopore Technologies, UK) according to manufacturer protocol with the following modifications: due to low PCR yield from low biomass input, the total volume from each PCR reaction was used as input, and CleanNGS SPRI beads were used for library clean-up steps. DNA end-prep reactions were scaled to accommodate higher volume input, and elution volume retained for direct use in later protocol steps. DNA concentration was measured using Qubit dsDNA HS Assay kit (Thermo Fisher Scientific, USA). Gel electrophoresis using Tapestation 2200 and D1000/High sensitivity D1000 screentapes (Agilent, USA) was used to validate product size and purity of a subset of amplicon libraries. The resulting sequencing libraries (2 x 114) were split into 4 pools, and each pool was loaded onto a MinION R10.4.1 flowcell and sequenced using the MinKNOW 22.12.7 software (Oxford Nanopore Technologies, UK). Data yield and quality were below that normally obtained from a MinION sequencing run due to the low DNA loaded, but sequencing nevertheless allowed to run until real-time data output was negligible. Reads were basecalled and demultiplexed with MinKNOW guppy g6.4.2 using the super accurate basecalling algorithm (config r10.4.1_400bps_sup.cfg) and custom barcodes.

### DNA extraction and sequencing from gut samples

Sample analysis was carried out by StarSEQ (Mainz, Germany) as follows: the samples were prepared using the Promega Maxwell CSC 48 robotic system with the Maxwell RSC Fecal Microbiome Kit. All samples were homogenized first using the Precellys Evolution beat beater (Bertin) with 4 x 20 s with a 30 s break between the steps at 7,800 rpm. The following primers were used for PCR amplification of bacterial rRNA gene variable region V4: 515F (5′-GTGNCAGCMGCCGCGGTAA-3′) and 806bR (5′-GGACTACNVGGGTWTCTAAT-3′). A one-step PCR system was used for the amplification ^64,65^. Sequencing was performed with the MiSeq system (Illumina) with 2 x 300 bp.

### Metabolomic analysis

Sample analysis was carried out by MS-Omics (Vedbæk, Denmark) using a Vanquish LC system coupled to Orbitrap Exploris 240 MS (Thermo Fisher Scientific). An electrospray ionization interface was used as ionization source. Analysis was performed in positive and negative ionization mode under polarity switching. The UPLC-MS measurements were performed using a slightly modified version of the protocol described previously ^67^. Peak areas were extracted using Compound Discoverer 3.2 (Thermo Fisher Scientific). Identification of compounds were performed at four levels: Level 1: identification by retention times (compared against in-house authentic standards), accurate mass (with an accepted deviation of 3 ppm), and MS/MS spectra; Level 2a: identification by retention times (compared against in-house authentic standards), and accurate mass (with an accepted deviation of 3 ppm); Level 2b: identification by accurate mass (with an accepted deviation of 3 ppm), and MS/MS spectra; Level 3: identification by accurate mass alone against the Human Metabolome Database (with an accepted deviation of 3ppm) ^68^.

### Bioinformatics and statistical analysis

For 16S rRNA gene abV4-C, Illumina sequencing produced paired-end reads that were processed using the LotuS2 (v2.23) pipeline ^69^. Sequences were clustered into OTUs using the Uparse algorithm ^70^, with other parameters set to the default options in LotuS2. All the OTUs with mitochondrial annotations were discarded. For 16S rRNA gene bV18-A of nanopore, the emu (v3.4.5) ^71^ with RDP database was used to obtain a species-level taxonomic abundance. In order to eliminate the microbiological contamination from the kits and surroundings, we carried out the following decontamination procedures: 1) Remove the spurious OTUs with an abundance less than 0.25 % ^72^ in at least two samples for lung tissue samples and at least one sample for gut tissue samples. 2) For lung tissue samples, eliminate the OTUs with 10 reads in at least one sample in the PCR kit negative control or DNA extraction kit negative control or environment blank control. 3) Discard the OTUs, which were annotated as contamination by R package decontam (set method to “prevalence”) by comparing the OTU prevalence in tissue samples and control samples ^73^. Alpha diversity was estimated using the Chao1 and Shannon indices, and the *p*-value was determined by the Wilcoxon rank-sum test. Beta diversity was estimated using weighted UniFrac and between-group *p*-values were obtained using PERMANOVA. DESeq2 (v1.38.3) was employed for differential abundance analysis of microbes, and the raw *p*-value was adjusted for multiple testing using the Benjamini-Hochberg procedure. Only genera with a relative abundance of more than 0.01 % and a prevalence of more than 10 % were considered for testing. The R package usedist (v0.4.0) was employed to estimate the inter-group beta diversity for the microbiome (weighted UniFrac distance) and metabolome (Euclidean distance), and the *p*-value was determined by the Wilcoxon rank-sum test.

For metabolomics, data within annotation levels 1, 2a, and 2b were used. Metabolites with a Descriptive Power (DP) less than or equal to 2.5 were filtered out, as were metabolites that were missing in more than 30 % of all samples. The abundance data of metabolites were transformed using log_2_. The Wilcoxon rank-sum test was used to test for differential abundance of metabolites, and the *p*-value was adjusted for multiple testing using the Benjamini-Hochberg procedure. The global correlation between lung microbiome, lung metabolome, gut microbiome, and plasma metabolome was illustrated using the Procrustes method. The correlation value was calculated using the RV2 method of the R package MatrixCorrelation ^74^, and the *p*-value was obtained using a permutation test. The mediation analysis was performed using the hdmed (v 1.0.0, HIMA method) package in R ^75,76^. The correlation network between genera and metabolites was based on Spearman’s correlation.

### DNA extraction and quantification of *L. murinus* and *A. fumigatus* DNA abundance in lung samples by qPCR

DNA was extracted with DNeasy Blood & Tissue Kit (Qiagen) with following modifications: 375 μl of mouse lung homogenate (∼25 mg tissue) were centrifuged to remove liquid, resuspended in 315 µl ATL buffer with 35 µl proteinase K and homogenized with Lysing Matrix E (MP Biomedicals) at 6.5 m/s for 30 s. Homogenates were centrifuged at 11,000 *g* at 4 °C for 7 min and incubated at 56 °C for 1 h. The lysate was centrifuged at 11,000 *g* at RT for 7 min, 200 µl of the supernatant was used for the DNA extraction. Prior to extraction, the lysate was treated with 2 µg/µl (w/v) RNAse A (Macherey-Nagel). Genomic DNA of *L. murinus* MAM1902 was extracted from *L. murinus* overnight culture grown on Columbia Blood Agar (CBA), gDNA of *A. fumigatus* A1160*^+^* was extracted from an overnight culture grown in *Aspergillus* minimal medium (AMM) ^77^. DNA extraction from *L. murinus* MAM1902 and *A. fumigatus* A1160*^+^* was conducted using the protocol described above with longer homogenization time (2 min for *L. murinus* and 1.5 min for *A. fumigatus*). DNA concentration was measured using a Qubit dsDNA BR Assay kit (Thermo Fisher Scientific). DNA quantification by qPCR was performed using SensiFAST Probe Direct SuperMix (Meridian Bioscience Inc.). For the quantification of *L. murinus* DNA, forward and reverse primers (400 nM) and fluorescent probe (200 nM) for a gene encoding a penicillin-binding protein were used (L.m.PBP_F 5’-AAGCTTGGCGCATCGTCATC-3’, L.m.PBP_R 5’-CAACCGTGCCGTTCAAACTG-3’, L.m.PBP_probe 5’-6-FAM/CAGGTGCAT/ZEN/AATCCATCAAAGGCTTAGCCGTAGAA/IBkFQ-3’). For the quantification of *A. fumigatus* DNA, forward and reverse primers (400 nM) and a fluorescent probe (100 nM) for a gene encoding tubulin beta-2 subunit were used (A.f.tub-b2_F 5’-GAGCCCTTTTCCGACCTGAT-3’, A.f.tub-b2_R 5’-GGAACTCCTCCCGGATCTTG-3’, A.f.tub-b2_probe 5’-6-FAM/CCAGATCAC /ZEN/CCATTCTCTGGGCGGCGGAAC/IBkFQ-3’). For the quantification of mouse DNA, forward and reverse primers (400 nM) and a fluorescent probe (100 nM) for a gene encoding beta-actin were used (b_actin_F 5’-AGCACAGCTTCTTTGCAGCTCC-3’, b_actin_R 5’-TGGTGTCCGTTCTGAGTGATCC-3’, b_actin_probe 5’-6-FAM/AGCGGGCCT/ZEN/ TCGCTCTCTCGTGGCTAGTA/IBkFQ-3’). PCR was performed with initial denaturation step at 95 °C for 3 min followed by 40 cycles of denaturation at 95 °C for 15 s and annealing/extension at 58 °C for 20 s. For the *L. murinus* and *A. fumigatus* DNA quantification, ∼1 µg of the lung tissue DNA was used for each reaction. DNA content was calculated based on the standard curves obtained for the mouse gDNA, *L. murinus* gDNA, and *A. fumigatus* gDNA. To compare DNA abundance among groups, the statistical analysis was performed using the Wilcoxon rank-sum test. The Spearman’s correlation was used to determine the abundance association between *A. fumigatus* DNA and *L. murinus* DNA, only samples with detected *L. murinus* DNA were taken for the analysis.

### Isolation of bacteria

Bacteria were isolated from lungs and tracheae of 28 male and female C57BL/6 mice, a commonly used inbred mouse strain, from four different providers: the animal facility of the BioInstrumente-Zentrum (BIZ, Jena, Germany), the Zentrale Tierexperimentelle Einrichtung (ZTE, University of Münster, Germany), Janvier Labs (Le Genest-Saint-Isle, Saint Berthevin, France) - C57BL/6J mouse substrain, and Charles River Laboratories (Sulzfeld, Germany) - C57BL/J6N mouse substrain. Mice were kept in specific and opportunistic pathogen-free (SOPF) conditions (BIZ Jena) or specific pathogen-free (SPF) conditions (ZTE University of Münster, Janvier Labs, and Charles River Laboratories). Mice from ZTE University of Münster, and Janvier Labs were housed at the animal facility of the Leibniz-HKI for 4 weeks, mice from Charles River Laboratories were housed at the Leibniz-HKI for 8 weeks before they were sacrificed. All animals were treated in accordance with European animal welfare regulations. and euthanized either under the legal provisions of § 4(3) of the German Animal Welfare Act or as control groups in another project (license no. HKI-19-006, 03-025/16). For the preparation of lung samples, the left lung or the upper lobe of the right lung was removed. For isolation of bacteria, lung and tracheal samples were homogenized in 0.5-1 ml DPBS (Gibco) in Lysing Matrix D tubes (MP Biomedicals) using a FastPrep homogenizer (MP Biomedicals) at 6 m/s for 2 x 40 s. The equal volume of DPBS was used as a negative control. Homogenized samples were plated on CBA, Tryptic Soy Agar (TSA), and Schaedler Agar (only for anaerobic cultivation) and incubated in aerobic, anaerobic or CO_2_-rich conditions (∼5 % (v/v)) at 37 °C for up to 6 days. Anaerobic conditions were obtained using an anaerobic atmosphere-generating system (Oxoid AnaeroGen), CO_2_-rich conditions were obtained with a CO_2_Gen (Oxoid) system. Bacteria with distinct colony morphology were used for pure culture establishment, identification and characterization.

### Microorganisms and culture conditions

*L. murinus*, *Lactobacillus taiwanensis*, *Streptococcus danieliae* and *Staphylococcus xylosus* were isolated from the lower respiratory tract of C57BL/6 mice from the animal facility of the BIZ. *L. murinus*, *L. taiwanensis*, and *S. danieliae* were cultured on CBA and *S. xylosus* was cultured on TSA at 37 °C aerobically or anaerobically. When indicated, the medium was supplemented with 2 µg/ml erythromycin. The following *A. fumigatus* strains were used in this study: CEA10 ^78^ and its derivatives A1160^+ 79^ and the Δ*pptA* mutant strain ^21^, ATCC 46645 ^80^ and its mutant derivatives Δ*sidA* ^81^ and Δ*freB*/Δ*sidA* ^82^*. A. fumigatus* was cultured in AMM and on AMM agar plates. When required, the medium was supplemented with 10 mM lysine, 1.5 mM FeSO_4_ or 10 µM of the siderophore triacetylfusarinine C (TAFC) ^21^.

### Measurement of lung epithelial cell damage

Murine T7 alveolar epithelial cells ^83^ were cultivated in Ham’s F-12K medium (Gibco) supplemented with 10 % (v/v) FCS and 0.5 % (v/v) Insulin-Transferrin-Selenium (Thermo Fisher Scientific). For the experiment, the cells were seeded at a density of 7 x 10^5^ cells/well and cultured overnight in a 24-well plate (TPP) with 1 ml of Dulbecco’s Modified Eagle medium (DMEM, Gibco) supplemented with 10 % (v/v) FCS, 1 % (v/v) GlutaMAX (Gibco) at 37 °C and 5 % (v/v) CO_2_. The medium was exchanged by 500 µl of DMEM (Gibco) without phenol red supplemented with 10 % (v/v) FCS, 1 % (v/v) GlutaMAX (Gibco). Cells were infected with 50 µl of bacterial suspension in DPBS (for *S. danieliae* and *S. xylosus*) or in water (for *L. taiwanensis* and *L. murinus*) with an MOI=5. As *S. danieliae* forms chains with on average 20 bacterial cells per chain, the required amount of *S. danieliae* CFUs was calculated as the required number of cells divided by 20. To detect spontaneous LDH activity, depending on the bacterial species, 50 µl DPBS or water were added to the cells. Experiments were performed in triplicates. Cells were incubated at 37 °C and 5 % (v/v) CO_2_ for 24 h. Cytotoxicity test was performed with CyQUANT LDH Cytotoxicity Assay Kit (Thermo Fisher Scientific) according to the instructions provided by the supplier.

### Generation of a GFP-producing *L. murinus* strain

The *L. murinus*/pTRKH3-ermGFP strain was obtained by transformation of *L. murinus* MAM1902 with the expression vector pTRKH3-ermGFP (addgene plasmid # 27169 ^84^). For transformation of *L. murinus*, 90 ml Man-Rogosa-Sharpe (MRS) medium in a 100 ml Erlenmeyer flask were inoculated with *L. murinus* harvested from CBA plates with a starting OD_600_ = 0.2. The culture was incubated at 37 °C, 75 rpm up to an OD_600_ = 1. The culture was centrifuged and washed three times with 20 ml ice-cold 0.3 M sucrose, 10 % (v/v) glycerol solution in water. The pellet was finally resuspended in 1 ml 0.5 M sucrose, 10 % (v/v) glycerol solution in water. 100 µl of the thus obtained bacterial suspension was mixed with 1 µl plasmid DNA solution containing 750 ng DNA. All procedures were performed on ice. Transformation was performed by electroporation in a 1 mm gap electroporation cuvette (Biozym) with an Eporator (Eppendorf) set to 2,500 V. Transformants were selected on CBA supplemented with 0.8 µg/ml erythromycin by anaerobic incubation for 2 days.

### Determination of internalization of *L. murinus* by lung epithelial cells

Murine T7 alveolar epithelial cells were cultivated in DMEM supplemented with 4 % (v/v) FCS, 1 % (v/v) GlutaMAX (Gibco), and 0.5 % (v/v) Insulin-Transferrin-Selenium (Thermo Fisher Scientific) at 37 °C and 5 % (v/v) CO_2_. For the experiment, cells were seeded at a density of 3.1 x 10^5^/well and cultured overnight in a 12-well plate (TPP) with 2.5 ml of DMEM supplemented with 4 % (v/v) FCS, 1 % (v/v) GlutaMAX (Gibco) at 37 °C and 5 % (v/v) CO_2_. Cells were infected with *L. murinus*/pTRKH3-ermGFP with an MOI = 5. Cultures were supplemented with 2 µg/ml erythromycin. Plates were centrifuged for 5 min at 500 *g* to synchronize the uptake. After 6 h, samples were washed with DPBS to remove bacteria that were not internalized. Cells were detached from the well by incubation with 0.3 ml TrplE enzyme (Gibco) for 5 min at 37 °C and 5 % (v/v) CO_2_. Then, 0.7 ml medium with 10 % (v/v) FCS were added and the cells were transferred to the microcentrifuge tube and kept on ice. Samples were centrifuged at 600 *g* for 2 min at 4 °C. After removing the supernatant, samples were fixed with 1 % (v/v) paraformaldehyde, washed twice with DPBS, resuspended in 50 µl DPBS, and stained with 2 µl of 0.5 mg/ml fresh concanavalin A AF647 (Invitrogen). For the compensation of signal spill-over, the following controls were prepared: one sample of T7 cells infected with *L. murinus* was not stained with concanavalin A AF647, and one sample of uninfected cells was stained with concanavalin A AF647. Imaging flow cytometry was conducted with ImageStream (Amnis) using Inspire Software (Amnis) with parameters: 642 nm – 0.5 mV, 488 nm – 50 mV. Internalization analysis was performed with Idias software (Amnis) using Internalization Wizard.

### Co-cultivation of L. murinus and A. fumigatus

For co-culture experiments, 1 x 10^5^ CFU/ml of *L. murinus* MAM1902 and 1 x 10^5^ conidia/ml of *A. fumigatus* CEA10 were inoculated in 1 ml medium in a 24-well plate. Cultivation was performed in Ham’s F-12K (Kaighn’s modification) medium (Gibco) at 37 °C and 5 % (v/v) CO_2_ or synthetic cystic fibrosis medium (SCFM2) ^85^ without mucin and lactic acid aerobically at 37 °C. *L. murinus* monoculture was used as a control. After 24 h, cultures were serially diluted and plated on TSA for counting CFUs. Plates were incubated anaerobically to prevent fungal growth. For the analysis of the effect of the siderophore TAFC on bacterial growth, 1 x 10^5^ CFU/ml of *L. murinus* were cultivated in 1 ml Ham’s F-12K medium supplemented with 10 µM TAFC in a 24-well plate. Cultivation was performed at 37 °C and 5 % (v/v) CO_2_ for 24 h. Bacterial growth was evaluated by counting CFUs and was compared to that observed with the *L. murinus* culture without TAFC supplementation.

To analyze a possible effect of Fe^3+^ reduction to Fe^2+^ on bacterial growth by FreB iron reductase located on the fungal surface, 1 x 10^5^ CFU/ml of *L. murinus* and 1 x 10^5^ conidia/ml of *A. fumigatus* strain Δ*freB*/Δ*sidA*, *A. fumigatus* wild type ATCC 46645 and its Δ*sidA* deletion mutant, were co-cultivated in 1 ml Ham’s F-12K medium supplemented with 10 M TAFC in a 24-well plate. Cultivation was performed at 37 °C and 5 % (v/v) CO_2_ for 24 h. Bacterial growth was evaluated by CFU quantification and data were compared to the results obtained with an *L. murinus* monoculture cultivated under the same conditions.

For the analysis of the effect of *A. fumigatus* natural products, whose biosynthesis is dependent on phosphopantetheinyl transferase (PptA), on bacterial growth, 1 x 10^5^ CFU/ml of *L. murinus* and 1 x 10^5^ conidia/ml of *A. fumigatus* Δ*pptA* mutant strain were cultivated in 1 ml Ham’s F-12K medium supplemented with 10 µM TAFC, 10 mM L-lysine in a 24-well plate. Cultivation was performed at 37 °C and 5 % (v/v) CO_2_ for 24 h. Bacterial growth was evaluated by quantification of CFUs and data were compared to the results obtained with an *L. murinus* monoculture cultivated under the same conditions.

To evaluate the dependence of *L. murinus* growth promotion by *A. fumigatus* on the culture volume, 1 x 10^5^ CFU/ml of *L. murinus* and 1 x 10^5^ conidia/ml of *A. fumigatus* were inoculated in a 96-well plate containing 50, 75, 100, 150, 200, 250, and 300 µl of Ham’s F-12K medium and cultivated for 24 h at 37 °C and 5 % (v/v) CO_2_. Bacterial growth was evaluated by quantification of CFUs.

To analyze growth of *L. murinus* in monoculture and co-culture with *A. fumigatus* over 24 h, *L. murinus* in monoculture and co-culture were inoculated in 9 wells in 96-well plates containing 200 µl Ham’s F-12K medium as described above. Bacterial growth was evaluated by quantification of CFUs at time points 0, 3, 6, 9, 12, 15, 18, 21, and 24 h. At each time point, 20-50 µl culture broth was sampled from the wells and serially diluted, if necessary.

### Oxygen concentration measurement

The oxygen concentration was measured with an OxoPlate (Presens) according to the supplier. Briefly, mono- and co-cultures of 1 x 10^5^ CFU/ml of *L. murinus* and 1 x 10^5^ conidia/ml of *A. fumigatus* were inoculated in a 96-well flat bottom OxoPlate with 200 µl Ham’s F-12K medium or SCFM2. As references, 400 µl/well oxygen-depleted water (10 mg/ml Na_2_SO_3_, freshly dissolved in water) and 200 µl/well oxygen-saturated water were used. Wells with oxygen-depleted water were sealed with adhesive foil. Measurement of Ham’s F-12K cultures was performed with a ClarioStar plate-reader (MBG Labtech) with incubation at 37 °C with 5 % (v/v) CO_2._. Measurement of SCFM2 cultures was performed with an Infinite PRO plate-reader (Tecan) at 37 °C. Samples were incubated for 60 min before the first measurement. Measurements were performed every hour for 24 h at 540 nm excitation, 650 nm emission for the indicator signal and 540 nm excitation, 590 nm emission for the reference signal.

### Growth of *L. murinus* with oxygen scavengers

1 x 10^5^ CFU/ml of *L. murinus* were inoculated in a 24-well plate with 1 ml Ham’s F-12K medium, Ham’s F-12K medium supplemented with freshly prepared 1.5 mM FeSO_4_ or 4 mM ascorbic acid. After an incubation for 24 h at 37 °C and 5 % (v/v) CO_2_, bacterial growth was evaluated by counting CFUs.

### Scanning electron microscopy

*L. murinus* in monoculture and co-culture with *A. fumigatus* were grown in a 24-well plate with 1 ml Ham’s F-12K medium 24 h at 37 °C and 5 % (v/v) CO_2_. The samples were washed with medium, transferred to poly-D-lysine coated glass cover slips, and fixed with freshly prepared modified Karnovsky fixative (4 % w/v paraformaldehyde, 2.5 % v/v glutaraldehyde in 0.1 M sodium cacodylate buffer, pH 7.4) for 1 h at room temperature. After washing 3 times for 15 min each with 0.1 M sodium cacodylate buffer (pH 7.4) the cells were post-fixed with 2 % (w/v) osmiumtetroxide for 1 h at room temperature. Subsequently, the samples were dehydrated in ascending ethanol concentrations (30, 50, 70, 90 and 100 %) for 15 min each. Next, the samples were critical-point dried using liquid CO_2_ and sputter coated with gold (thickness approx. 2 nm) using a CCU-010 sputter coater (safematic GmbH, Zizers, Switzerland). Finally, the specimens were investigated with a field emission SEM LEO-1530 Gemini (Carl Zeiss NTS GmbH, Oberkochen, Germany).

