## Supplementary Figures S1-S4 for "The murine lung microbiome is dysbalanced by the human pathogenic fungus *Aspergillus fumigatus* resulting in enrichment of anaerobic bacteria"

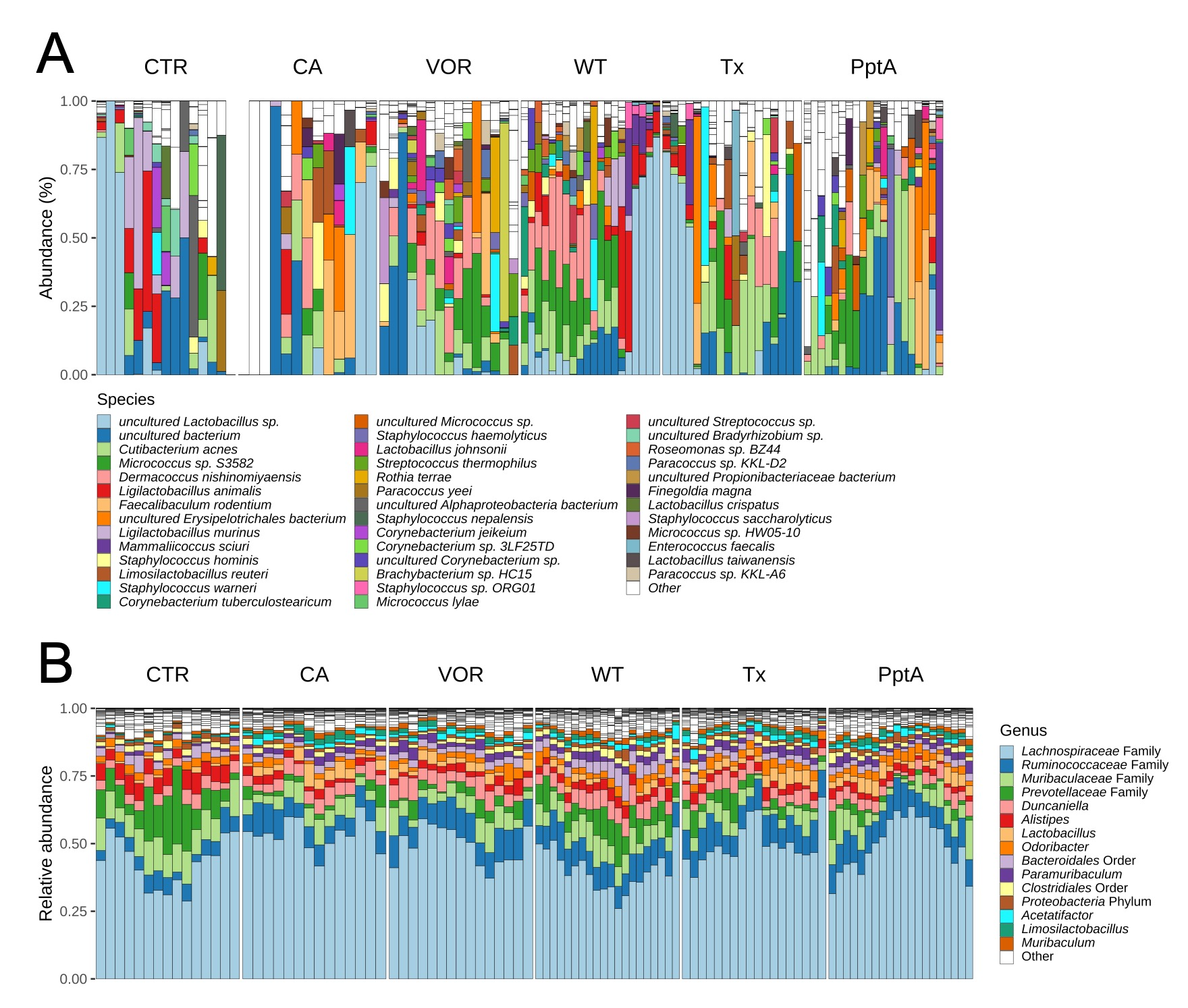

**Figure S1.** The composition of the lung and gut microbiome changes dependent on the treatment of mice. A) Composition plot for the lung microbiome based on 16S rDNA with identification at the species level. B) Composition plot for the lung microbiome based on 16S rDNA. Abbreviations: CTR, control mice; CA, cortisone acetate-immunosuppressed mice; VOR, cortisone acetate-immunosuppressed mice treated with voriconazole for 3 days; WT, cortisone acetate-immunosuppressed mice infected with *A. fumigatus*; Tx, mice immunosuppressed with cortisone acetate, infected with *A. fumigatus* and treated with voriconazole; PptA, mice immunosuppressed with cortisone acetate and infected with conidia of the avirulent *A. fumigatus* ∆*pptA* strain. Each bar represents data from a single animal.

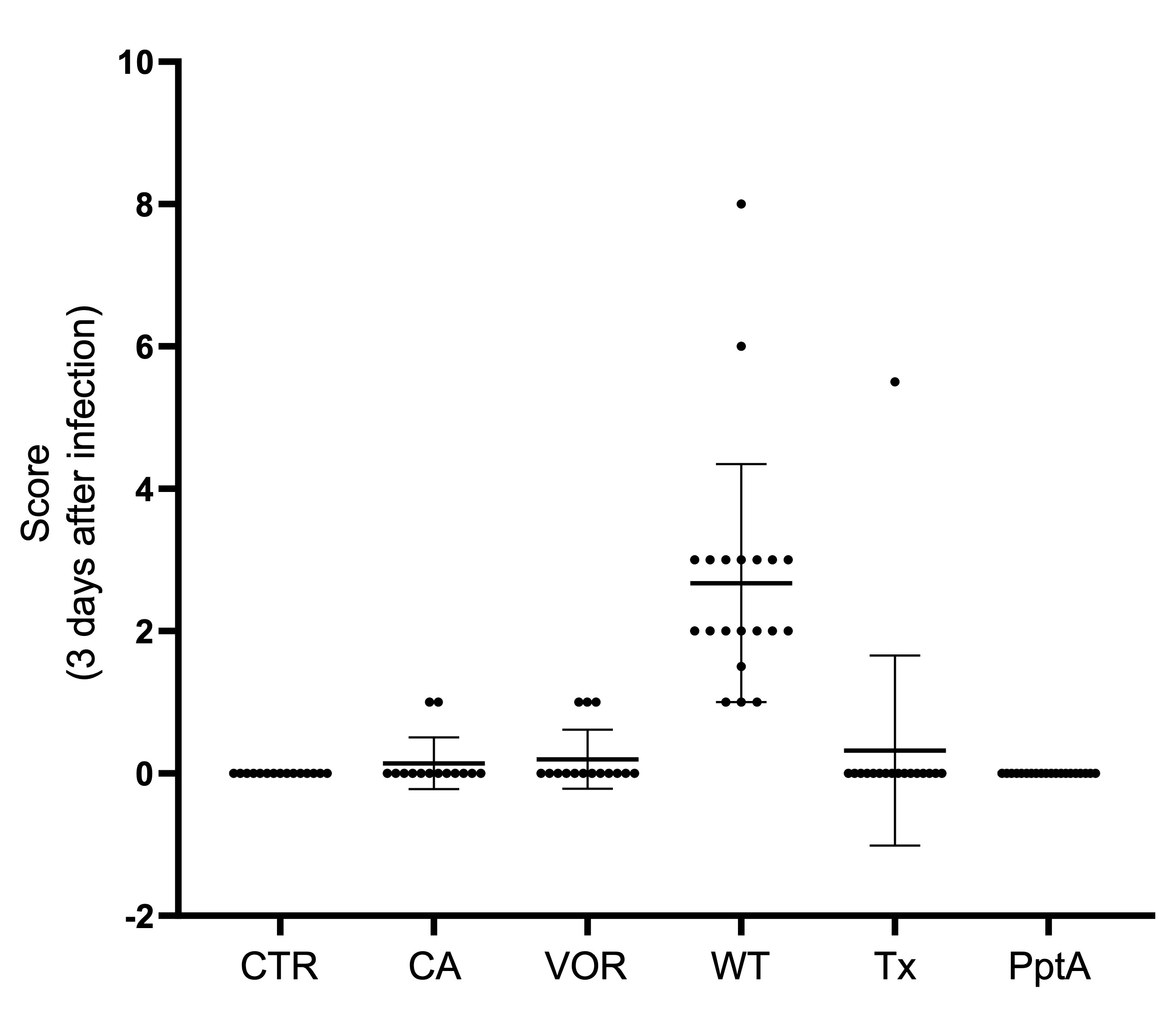

**Figure S2.** Clinical symptoms scores on day 3 after infection. Immunocompromised mice infected with *A. fumigatus* wild type (WT) showed the expected symptoms of infection on the day of sacrifice. Abbreviations: control mice (CTR); cortisone acetate-immunosuppressed mice (CA); cortisone acetate-immunosuppressed mice treated with voriconazole for 3 days (VOR); cortisone acetate-immunosuppressed mice infected with *A. fumigatus* wild-type conidia (WT); mice immunosuppressed with cortisone acetate, infected with *A. fumigatus* and treated with voriconazole (Tx); and mice immunosuppressed with cortisone acetate and infected with conidia of the avirulent *A. fumigatus* ∆*pptA* strain (PptA).

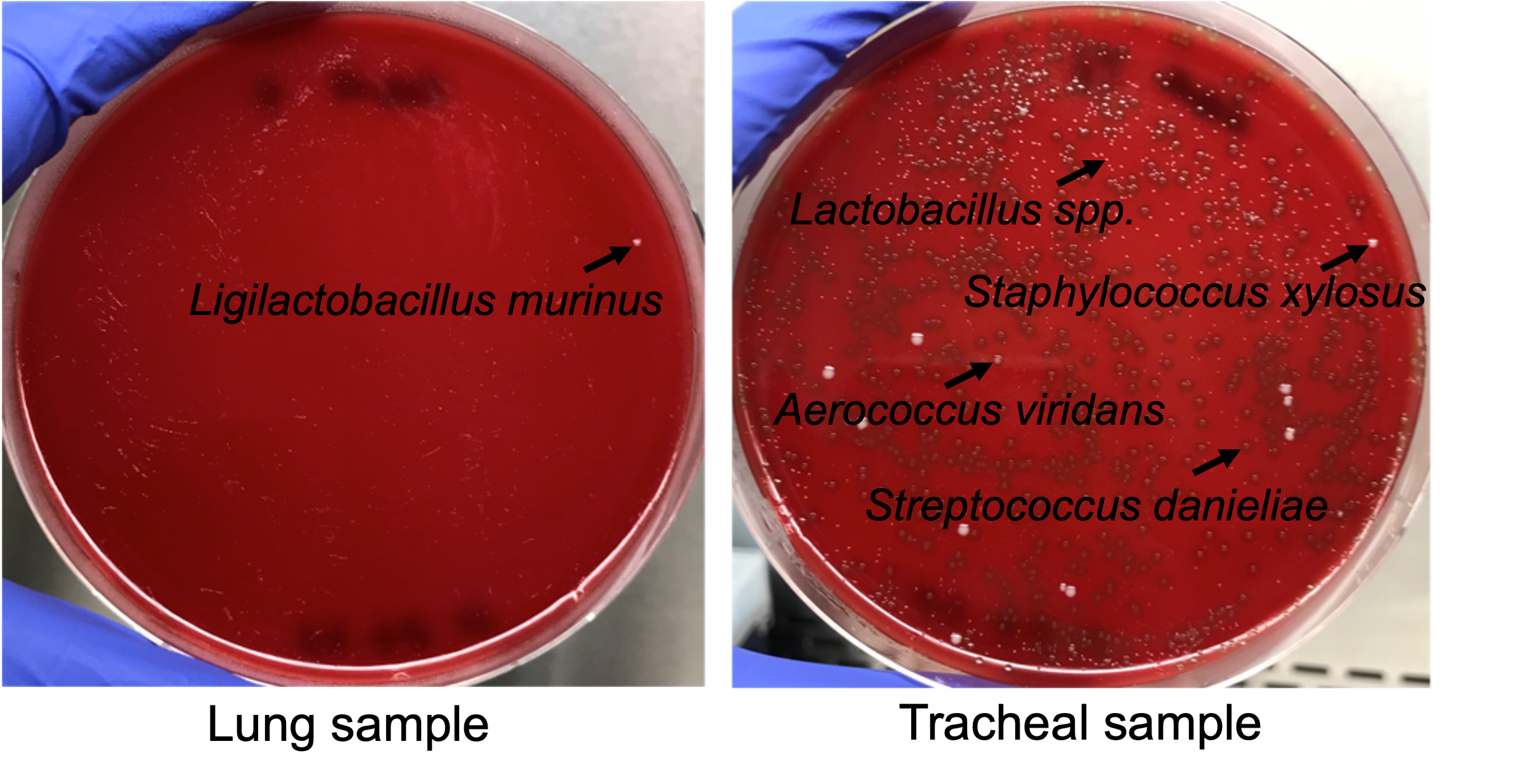

**Figure S3.** Colonies grown from isolated bacteria from lung and tracheal samples were cultivated on Columbia Blood Agar plates.

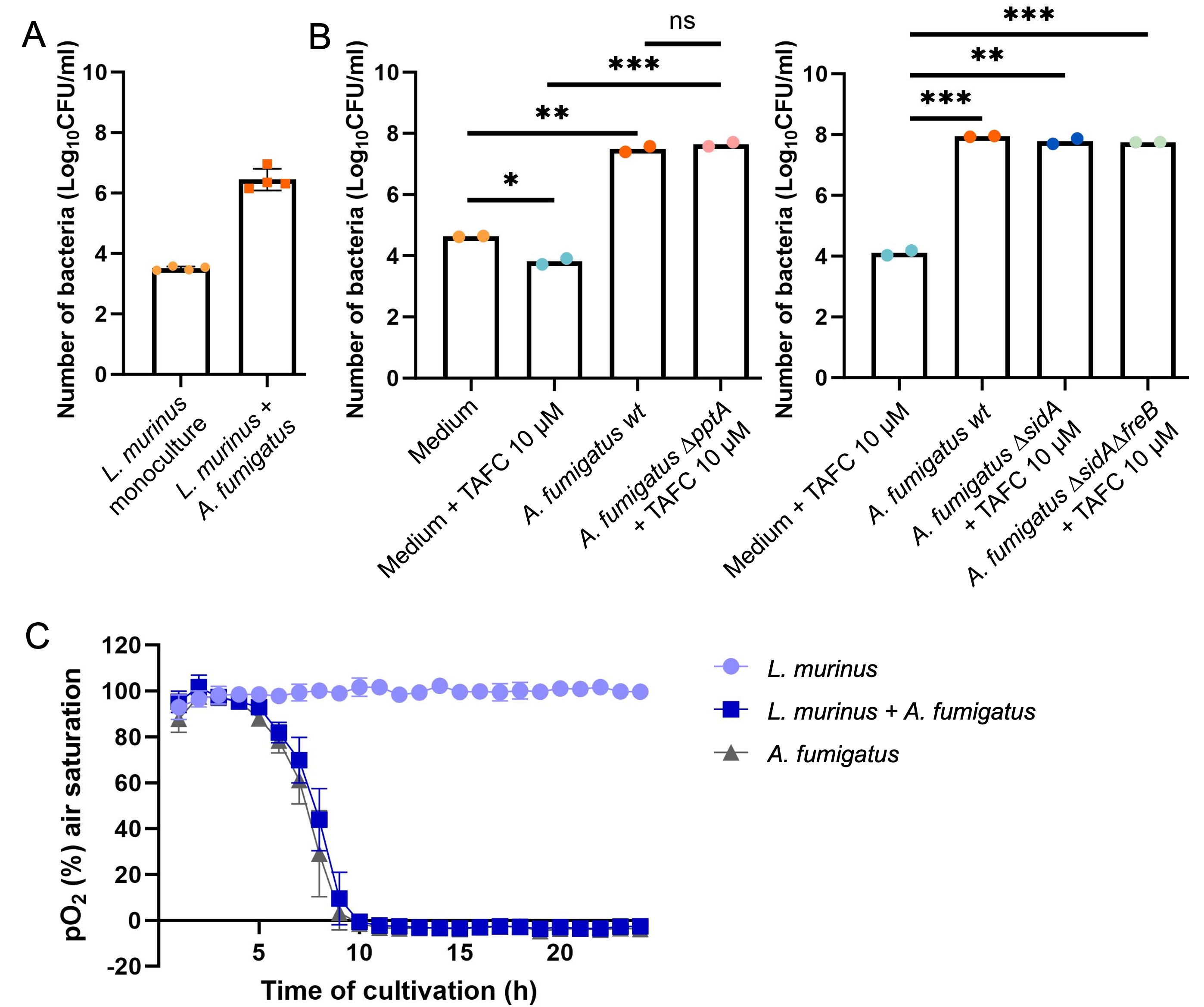
 **Figure S4.** *A. fumigatus* promotes the growth of *L. murinus* by creating a microaerophilic environment. A) Number of CFUs of *L. murinus* monoculture and co-culture with *A. fumigatus* after 24 hours of cultivation in modified SCFM2. B) *L. murinus* growth promotion by *A. fumigatus* is independent of TAFC siderophore production (∆*sidA*), the production of other natural products (∆*pptA*) or the reduction of Fe^3+^ to Fe^2+^ by the surface iron reductase FreB (∆*sidA*∆*freB*). Number of CFUs of *L. murinus* in monoculture and co-culture with different *A. fumigatus* strains. Cultivation was carried out in Ham’s F12-K medium with or without supplementation with 10 µM TAFC siderophore in 24-well plates for 24 h. C) pO_2_ measurement for *L. murinus* and *A. fumigatus* monocultures and co-culture of *L. murinus* and *A. fumigatus* over 24 h in modified SCFM2.
