## Supplementary Tables S1-S4 for "The murine lung microbiome is dysbalanced by the human pathogenic fungus *Aspergillus fumigatus* resulting in enrichment of anaerobic bacteria"

**Table S1.** Lung genera with differential abundance (by *p* < 0.05 and |log_2_FC| > log_2_(1.5); DESeq2).

| Comparison | OTU | Fold change | log_2_FC | *p* value |
| --- | --- | --- | --- | --- |
| CTR vs CA | *Bifidobacterium* | 6788630.729 | 22.695 | 1.183E-14 |
| CTR vs CA | *Faecalibaculum* | 268.091 | 8.067 | 9.222E-05 |
| CTR vs CA | *Staphylococcaceae* Family | 62.712 | 5.971 | 0.041 |
| CTR vs CA | *Anaerococcus* | 52.958 | 5.727 | 0.012 |
| CTR vs CA | *Methanothermobacter* | 33.072 | 5.048 | 0.002 |
| CTR vs CA | *Lactobacillus* | 27.968 | 4.806 | 0.039 |
| CTR vs CA | *Photobacterium* | 0.124 | -3.015 | 0.045 |
| CTR vs CA | *Latilactobacillus* | 0.031 | -4.999 | 0.016 |
| CA vs VOR | *Buchnera* | 25430808.076 | 24.600 | 5.798E-17 |
| CA vs VOR | *Cyanobacteria* Class | 59.673 | 5.899 | 0.044 |
| CA vs VOR | *Pasteurellaceae* Family | 30.501 | 4.931 | 0.025 |
| CA vs VOR | *Bacteroidetes* Phylum | 30.337 | 4.923 | 0.043 |
| CA vs VOR | *Faecalibaculum* | 0.038 | -4.711 | 0.011 |
| CA vs VOR | *Bifidobacterium* | 1.592E-07 | -22.583 | 1.533E-14 |
| CA vs WT | *Lactobacillales* Order | 281.139 | 8.135 | 0.006 |
| CA vs WT | *Paracoccus* | 242.403 | 7.921 | 2.678E-04 |
| CA vs WT | *Staphylococcaceae* Family | 219.361 | 7.777 | 9.576E-06 |
| CA vs WT | *Bacillales* Order | 60.282 | 5.914 | 0.023 |
| CA vs WT | *Chryseobacterium* | 34.371 | 5.103 | 0.042 |
| CA vs WT | *Corynebacterium* | 22.375 | 4.484 | 0.008 |
| CA vs WT | *Ligilactobacillus* | 7.133 | 2.835 | 0.023 |
| CA vs WT | *Lactobacillus* | 0.004 | -8.098 | 1.138E-05 |
| CA vs WT | *Photobacterium* | 0.002 | -8.649 | 4.387E-07 |
| CA vs WT | *Faecalibaculum* | 0.002 | -9.265 | 1.701E-05 |
| CA vs PptA | *Schaedlerella* | 8481819.462 | 23.016 | 1.250E-14 |
| CA vs PptA | *Paracoccus* | 234.978 | 7.876 | 3.307E-04 |
| CA vs PptA | *Methanothermobacter* | 0.051 | -4.299 | 0.006 |
| CA vs PptA | *Bradyrhizobium* | 0.016 | -5.940 | 0.003 |
| CA vs PptA | *Photobacterium* | 0.002 | -8.676 | 3.887E-05 |
| WT vs Tx | *Brachybacterium* | 463.446 | 8.856 | 0.0002 |
| WT vs Tx | *Faecalibaculum* | 187.274 | 7.549 | 0.0002 |
| WT vs Tx | *Limosilactobacillus* | 60.783 | 5.926 | 3.837E-05 |
| WT vs Tx | *Anaerococcus* | 0.118 | -3.085 | 0.046 |
| WT vs Tx | *Chryseobacterium* | 0.049 | -4.337 | 0.018 |
| WT vs Tx | *Staphylococcaceae* Family | 0.033 | -4.926 | 0.002 |
| WT vs Tx | *Bradyrhizobium* | 0.027 | -5.192 | 0.001 |
| WT vs Tx | *Enterococcus* | 0.017 | -5.918 | 0.017 |
| WT vs Tx | *Bacillales* Order | 0.016 | -5.932 | 0.020 |
| WT vs Tx | *Pasteurellaceae* Family | 0.013 | -6.232 | 0.005 |
| WT vs Tx | *Lactobacillales* Order | 0.005 | -7.573 | 0.001 |
| WT vs Tx | *Methanothermobacter* | 0.002 | -9.266 | 1.536E-06 |

**Table S2.** Gut genera with differential abundance (by *p* < 0.05 and |log_2_FC| > log_2_(1.5); DESeq2).

| Comparison | OTU | Fold change | log2FC | p value |
| --- | --- | --- | --- | --- |
| CTR vs CA | *Turicibacter* | 720.263 | 9.492 | 4.000E-16 |
| CTR vs CA | *Faecalibaculum* | 46.022 | 5.524 | 1.661E-22 |
| CTR vs CA | *Olsenella* | 7.092 | 2.826 | 1.552E-07 |
| CTR vs CA | *Staphylococcus* | 6.787 | 2.763 | 0.017 |
| CTR vs CA | *Erysipelotrichaceae* Family | 6.077 | 2.603 | 1.146E-04 |
| CTR vs CA | *Limosilactobacillus* | 6.029 | 2.592 | 7.064E-12 |
| CTR vs CA | *Lactobacillus* | 4.397 | 2.137 | 1.978E-07 |
| CTR vs CA | *Acetatifactor* | 2.737 | 1.452 | 1.422E-09 |
| CTR vs CA | *Paramuribaculum* | 2.671 | 1.418 | 3.157E-11 |
| CTR vs CA | *Desulfovibrionaceae* Family | 2.562 | 1.357 | 4.196E-07 |
| CTR vs CA | *Muribaculum* | 1.669 | 0.739 | 3.478E-04 |
| CTR vs CA | *Anaerotignum* | 0.658 | -0.603 | 0.038 |
| CTR vs CA | *Bacteroidales* Order | 0.658 | -0.605 | 1.249E-04 |
| CTR vs CA | *Muribaculaceae* Family | 0.611 | -0.711 | 0.002 |
| CTR vs CA | *Firmicutes* Phylum | 0.549 | -0.864 | 0.021 |
| CTR vs CA | *Proteobacteria* Phylum | 0.498 | -1.007 | 2.493E-09 |
| CTR vs CA | *Alistipes* | 0.411 | -1.282 | 3.905E-16 |
| CTR vs CA | *Dorea* | 0.409 | -1.289 | 0.002 |
| CTR vs CA | *Neglecta* | 0.363 | -1.463 | 6.475E-09 |
| CTR vs CA | *Clostridium XlVa* | 0.305 | -1.714 | 9.757E-24 |
| CTR vs CA | *Bacteroides* | 0.207 | -2.271 | 1.048E-14 |
| CTR vs CA | *Prevotellaceae* Family | 0.207 | -2.274 | 1.739E-10 |
| CTR vs CA | *Eisenbergiella* | 0.115 | -3.116 | 0.021 |
| CA vs VOR | *Staphylococcus* | 5.742 | 2.522 | 3.348E-04 |
| CA vs VOR | *Bacteroidetes* Phylum | 1.941 | 0.957 | 0.001 |
| CA vs VOR | *Bacteroides* | 1.772 | 0.826 | 0.007 |
| CA vs VOR | *Dorea* | 0.636 | -0.653 | 0.012 |
| CA vs VOR | *Akkermansia* | 0.016 | -5.946 | 0.004 |
| CA vs WT | *Escherichia/Shigella* | 20.036 | 4.325 | 0.007 |
| CA vs WT | *Bacteroidetes* Phylum | 3.355 | 1.746 | 4.497E-08 |
| CA vs WT | *Staphylococcus* | 3.312 | 1.728 | 0.014 |
| CA vs WT | *Bacteroides* | 2.645 | 1.404 | 4.223E-06 |
| CA vs WT | *Prevotellaceae* Family | 2.563 | 1.358 | 6.681E-06 |
| CA vs WT | *Dysosmobacter* | 2.520 | 1.333 | 1.432E-06 |
| CA vs WT | *Clostridia* Class | 2.255 | 1.173 | 0.024 |
| CA vs WT | *Bacteroidales* Order | 1.955 | 0.967 | 1.407E-09 |
| CA vs WT | *Lachnospiraceae* Family | 0.653 | -0.616 | 2.517E-05 |
| CA vs WT | *Schaedlerella* | 0.626 | -0.675 | 0.019 |
| CA vs WT | *Olsenella* | 0.447 | -1.160 | 0.013 |
| CA vs WT | *Lactobacillus* | 0.408 | -1.294 | 0.003 |
| CA vs WT | *Limosilactobacillus* | 0.399 | -1.326 | 0.002 |
| CA vs PptA | *Staphylococcus* | 2.976 | 1.574 | 0.030 |
| CA vs PptA | *Proteobacteria* Phylum | 1.638 | 0.712 | 2.747E-04 |
| CA vs PptA | *Dysosmobacter* | 1.580 | 0.660 | 0.018 |
| CA vs PptA | *Limosilactobacillus* | 0.615 | -0.702 | 0.021 |
| CA vs PptA | *Ligilactobacillus* | 0.363 | -1.461 | 7.937E-09 |
| CA vs PptA | *Turicibacter* | 0.013 | -6.255 | 2.742E-12 |
| WT vs Tx | *Lactobacillus* | 2.198 | 1.136 | 0.007 |
| WT vs Tx | *Limosilactobacillus* | 1.736 | 0.796 | 0.042 |
| WT vs Tx | *Phocea* | 1.693 | 0.760 | 0.030 |
| WT vs Tx | *Bacteroidales* Order | 0.626 | -0.676 | 1.603E-05 |
| WT vs Tx | *Prevotellaceae* Family | 0.593 | -0.753 | 0.014 |
| WT vs Tx | *Staphylococcus* | 0.361 | -1.468 | 0.022 |
| WT vs Tx | *Clostridia* Class | 0.295 | -1.760 | 8.288E-04 |

**Table S3.** Lung metabolites with differential abundance (by *p* < 0.05 and |log_2_FC| > log_2_(1.5); DESeq2).

| Comparison | Metabolite | Fold change | log_2_FC | *p* value | Annotation level |
| --- | --- | --- | --- | --- | --- |
| CTR vs CA | Phosphocholine | 2.701 | 1.433 | 4.210E-04 | 1 |
| CTR vs CA | Butyrylcarnitine | 2.646 | 1.404 | 7.740E-07 | 2a |
| CTR vs CA | Isovalerylcarnitine | 2.510 | 1.328 | 3.090E-07 | 1 |
| CTR vs CA | D-Glucose 6-phosphate | 2.128 | 1.090 | 6.290E-04 | 2b |
| CTR vs CA | Propionylcarnitine | 2.098 | 1.069 | 0.005 | 1 |
| CTR vs CA | Dehydronorketamine | 2.079 | 1.056 | 1.110E-03 | 2b |
| CTR vs CA | Glycerol 3-phosphate | 1.800 | 0.848 | 7.010E-06 | 2b |
| CTR vs CA | 3-Hydroxybutyric acid | 1.576 | 0.656 | 0.003 | 1 |
| CTR vs CA | Ornithine | 0.655 | -0.610 | 9.250E-04 | 1 |
| CTR vs CA | Betaine | 0.628 | -0.672 | 1.110E-04 | 2a |
| CTR vs CA | Guanosine | 0.614 | -0.703 | 5.160E-04 | 2a |
| CTR vs CA | Homoarginine | 0.614 | -0.705 | 1.600E-03 | 2a |
| CTR vs CA | Acetylcholine | 0.581 | -0.783 | 4.060E-05 | 2b |
| CTR vs CA | Glutamic acid | 0.577 | -0.793 | 5.030E-06 | 2a |
| CTR vs CA | 2-Deoxyribose 5-phosphate | 0.561 | -0.834 | 2.760E-04 | 2b |
| CTR vs CA | γ-Aminobutyric acid | 0.552 | -0.858 | 9.620E-06 | 2a |
| CTR vs CA | O-Acetylserine | 0.544 | -0.878 | 1.310E-05 | 2a |
| CTR vs CA | Trigonelline | 0.536 | -0.900 | 2.500E-06 | 1 |
| CTR vs CA | Cytidine | 0.492 | -1.022 | 7.740E-07 | 1 |
| CTR vs CA | Xanthine | 0.479 | -1.061 | 2.350E-05 | 1 |
| CTR vs CA | dUridine | 0.456 | -1.133 | 3.100E-05 | 2a |
| CTR vs CA | Trimethylamine N-Oxide | 0.444 | -1.172 | 4.060E-05 | 2a |
| CTR vs CA | Xanthosine | 0.430 | -1.218 | 3.580E-06 | 2a |
| CTR vs CA | 1-Methyl Adenosine | 0.385 | -1.377 | 4.900E-07 | 2a |
| CA vs WT | Methionine sulfoxide | 2.562 | 1.358 | 2.870E-09 | 2a |
| CA vs WT | 1-Methylhistidine | 1.755 | 0.811 | 5.360E-07 | 2b |
| CA vs WT | Uridine | 1.519 | 0.603 | 0.003 | 1 |
| CA vs WT | Isovalerylcarnitine | 0.664 | -0.591 | 0.006 | 1 |
| CA vs WT | Butyrylcarnitine | 0.555 | -0.850 | 1.270E-05 | 2a |
| CA vs WT | Trigonelline | 0.393 | -1.349 | 3.720E-05 | 1 |
| CA vs WT | VLH | 0.284 | -1.818 | 6.290E-04 | 2b |
| CA vs PptA | Dehydronorketamine | 16.387 | 4.035 | 1.440E-09 | 2b |
| CA vs PptA | dUridine | 6.119 | 2.613 | 6.470E-08 | 2b |
| CA vs PptA | Uridine | 3.019 | 1.594 | 9.630E-08 | 1 |
| CA vs PptA | Clomazone | 2.820 | 1.496 | 1.010E-05 | 2b |
| CA vs PptA | Methionine sulfoxide | 2.650 | 1.406 | 1.440E-09 | 2a |
| CA vs PptA | 1-Methyl Adenosine | 2.362 | 1.240 | 7.300E-07 | 2a |
| CA vs PptA | Xanthine | 1.916 | 0.938 | 6.300E-06 | 1 |
| CA vs PptA | Thymidine | 1.915 | 0.937 | 9.810E-07 | 2a |
| CA vs PptA | Prostaglandin E2 | 1.846 | 0.884 | 1.010E-08 | 2a |
| CA vs PptA | Thymine | 1.763 | 0.818 | 1.310E-06 | 1 |
| CA vs PptA | dCytidine | 1.739 | 0.798 | 2.000E-07 | 1 |
| CA vs PptA | Lysine | 1.731 | 0.791 | 1.270E-05 | 1 |
| CA vs PptA | Trigonelline | 1.601 | 0.679 | 2.450E-05 | 1 |
| CA vs PptA | Creatinine | 1.596 | 0.675 | 2.270E-06 | 2a |
| CA vs PptA | Ornithine | 1.594 | 0.673 | 3.720E-05 | 1 |
| CA vs PptA | Phenylalanine | 1.592 | 0.671 | 3.910E-07 | 2a |
| CA vs PptA | Tyrosine | 1.574 | 0.654 | 7.300E-07 | 1 |
| CA vs PptA | O-Acetylserine | 1.545 | 0.628 | 3.720E-05 | 2a |
| CA vs PptA | Arginine | 1.537 | 0.620 | 1.980E-05 | 1 |
| CA vs PptA | N3,N4-Dimethyl-L-arginine | 1.531 | 0.615 | 5.550E-05 | 2b |
| CA vs PptA | Cytidine | 1.530 | 0.613 | 1.310E-06 | 1 |
| CA vs PptA | Glycerol 3-phosphate | 1.528 | 0.611 | 1.590E-05 | 2b |
| CA vs PptA | γ-Aminobutyric acid | 1.525 | 0.609 | 1.010E-05 | 2a |
| CA vs PptA | Pyroglutamic acid | 1.501 | 0.586 | 1.390E-07 | 2a |
| CA vs PptA | Trimethylamine N-Oxide | 0.618 | -0.694 | 0.015 | 2a |
| CA vs PptA | 3-Hydroxybutyric acid | 0.594 | -0.751 | 0.014 | 1 |
| CA vs PptA | Hypoxanthine | 0.566 | -0.822 | 0.015 | 1 |
| CA vs PptA | Inosine | 0.476 | -1.071 | 0.019 | 2a |
| CA vs PptA | Allopurinol | 0.428 | -1.224 | 5.390E-04 | 1 |
| CA vs PptA | dInosine | 0.348 | -1.523 | 4.610E-04 | 2a |
| CA vs PptA | 10-(Hydroxymethyl)-4-methyl-8,11-dioxa-2,6-diazatricyclo[7.2.1.02,7]dodeca-3,6-dien-5-one | 0.311 | -1.685 | 6.470E-08 | 2b |
| CA vs PptA | Cystine | 0.114 | -3.136 | 3.540E-06 | 2a |
| CA vs PptA | 2-Deoxyribose 5-phosphate | 0.013 | -6.316 | 6.850E-07 | 2b |
| WT vs Tx | Trigonelline | 2.612 | 1.3845 | 6.360E-06 | 1 |
| WT vs Tx | Butyrylcarnitine | 1.738 | 0.797 | 1.810E-06 | 2a |
| WT vs Tx | 2-Deoxyribose 5-phosphate | 1.522 | 0.606 | 0.011 | 2b |
| WT vs Tx | 3-Hydroxybutyric acid | 0.541 | -0.886 | 0.025 | 1 |
| WT vs Tx | Methionine sulfoxide | 0.535 | -0.901 | 5.550E-05 | 2a |

**Table S4.** Plasma metabolites with differential abundance (by *p* < 0.05 and |log_2_FC| > log_2_(1.5); DESeq2).

| Comparison | Metabolite | Fold change | log_2_FC | *p* value | Annotation level |
| --- | --- | --- | --- | --- | --- |
| CTR vs CA | 20 β-Dihydroprednisolone | 82.906 | 6.373 | 4.640E-06 | 2b |
| CTR vs CA | Cortisone | 75.634 | 6.241 | 2.580E-08 | 1 |
| CTR vs CA | Hyocholic acid | 56.873 | 5.830 | 4.800E-05 | 2a |
| CTR vs CA | Cholic acid | 54.577 | 5.770 | 2.760E-04 | 1 |
| CTR vs CA | a/b/ω-Muricholic acid | 35.072 | 5.132 | 2.760E-04 | 2a |
| CTR vs CA | 5,6-dimethyl-4-oxo-4H-pyran-2-carboxylic acid | 23.190 | 4.535 | 2.580E-08 | 2b |
| CTR vs CA | 3,5-Dimethoxybenzoic acid | 11.631 | 3.540 | 2.580E-08 | 2b |
| CTR vs CA | Dehydronorketamine_3 | 7.177 | 2.843 | 2.580E-08 | 2b |
| CTR vs CA | Decanoylcarnitine | 5.046 | 2.335 | 5.160E-08 | 1 |
| CTR vs CA | 5-Methoxysalicylic acid | 4.392 | 2.135 | 1.410E-04 | 2a |
| CTR vs CA | Octanoylcarnitine | 4.293 | 2.102 | 2.580E-08 | 2a |
| CTR vs CA | 2-Hydroxyvaleric acid | 4.282 | 2.098 | 2.580E-08 | 2b |
| CTR vs CA | [(1S,4S,6S)-6-Isopropyl-3-methyl-4-{2-oxo-2-[4-(2-pyrimidinyl)-1-piperazinyl]ethyl}-2-cyclohexen-1-yl]acetonitrile | 4.095 | 2.034 | 8.730E-05 | 2b |
| CTR vs CA | 2-Hydroxyisocaproic acid | 3.691 | 1.884 | 2.580E-08 | 2a |
| CTR vs CA | Dihydroxybenzoic | 3.675 | 1.878 | 2.500E-06 | 1 |
| CTR vs CA | Cytosine | 3.613 | 1.853 | 5.030E-06 | 1 |
| CTR vs CA | 3,4-Dimethoxyphenylacetic acid | 3.347 | 1.743 | 5.030E-06 | 1 |
| CTR vs CA | 10-HDA | 3.285 | 1.716 | 2.350E-05 | 2b |
| CTR vs CA | N-Acetyl-cysteine | 3.248 | 1.700 | 0.023 | 1 |
| CTR vs CA | Dehydronorketamine_2 | 3.086 | 1.626 | 9.190E-06 | 2b |
| CTR vs CA | Norketamine_1 | 3.081 | 1.624 | 2.500E-06 | 2b |
| CTR vs CA | β-D-Glucopyranuronic acid | 3.036 | 1.602 | 3.090E-07 | 2b |
| CTR vs CA | Salicylic acid | 2.953 | 1.562 | 4.900E-07 | 2a |
| CTR vs CA | Isovalerylcarnitine | 2.750 | 1.459 | 3.090E-07 | 1 |
| CTR vs CA | Glucuronic acid-3,6-lactone | 2.675 | 1.420 | 5.160E-08 | 2b |
| CTR vs CA | 5-Methylcytosine | 2.597 | 1.377 | 1.110E-03 | 2a |
| CTR vs CA | Dehydronorketamine_1 | 2.560 | 1.356 | 8.730E-05 | 2b |
| CTR vs CA | Glucuronic acid | 2.539 | 1.344 | 2.210E-04 | 2a |
| CTR vs CA | Norketamine_2 | 2.498 | 1.321 | 2.500E-06 | 2b |
| CTR vs CA | 3-Phenyllactic acid | 2.485 | 1.313 | 9.620E-06 | 1 |
| CTR vs CA | 3-(propan-2-yl)-octahydropyrrolo[1,2-a]pyrazine-1,4-dione_2 | 2.399 | 1.263 | 0.011 | 2b |
| CTR vs CA | 2-Hydroxyoctanoic acid | 2.289 | 1.195 | 1.770E-04 | 1 |
| CTR vs CA | Indole-3-lactic acid | 2.231 | 1.158 | 2.500E-06 | 2a |
| CTR vs CA | 3-Hydroxybutyric acid | 2.226 | 1.155 | 9.250E-04 | 1 |
| CTR vs CA | Pyrene | 2.199 | 1.137 | 1.160E-06 | 2b |
| CTR vs CA | 3-Hydroxy-2-methyl-4-pyrone | 2.098 | 1.069 | 1.110E-03 | 2a |
| CTR vs CA | (R)-Equol | 1.937 | 0.954 | 0.016 | 2b |
| CTR vs CA | Crotonic acid_2 | 1.805 | 0.852 | 4.210E-04 | 2b |
| CTR vs CA | Butyrylcarnitine | 1.795 | 0.844 | 4.900E-07 | 2a |
| CTR vs CA | Glycine anhydride | 1.762 | 0.817 | 1.110E-03 | 2b |
| CTR vs CA | Indole-3-acrylic acid | 1.700 | 0.766 | 0.004 | 2a |
| CTR vs CA | 4-Hydroxybenzaldehyde | 1.690 | 0.757 | 0.011 | 2a |
| CTR vs CA | Pyruvic acid | 1.670 | 0.740 | 2.500E-06 | 1 |
| CTR vs CA | Hexanoylcarnitine | 1.659 | 0.731 | 9.250E-04 | 2b |
| CTR vs CA | 3-Methylhistidine | 1.657 | 0.729 | 3.090E-07 | 1 |
| CTR vs CA | Rexamino | 1.634 | 0.709 | 0.022 | 2b |
| CTR vs CA | Traumatic acid | 1.624 | 0.699 | 4.210E-04 | 2a |
| CTR vs CA | Threonine | 1.574 | 0.654 | 4.900E-07 | 2a |
| CTR vs CA | 3-Ureidopropionic acid | 1.569 | 0.650 | 0.008 | 2a |
| CTR vs CA | Homocitrulline | 1.565 | 0.647 | 2.350E-05 | 2a |
| CTR vs CA | Glycine | 1.556 | 0.638 | 1.810E-07 | 1 |
| CTR vs CA | Prolylglycine | 1.509 | 0.593 | 6.820E-05 | 2b |
| CTR vs CA | Glyoxylic acid | 0.663 | -0.593 | 0.037 | 2a |
| CTR vs CA | Spermine | 0.658 | -0.603 | 9.250E-04 | 2a |
| CTR vs CA | N-Acetyl-lysine | 0.656 | -0.608 | 9.250E-04 | 1 |
| CTR vs CA | Isobutyric acid_2 | 0.653 | -0.615 | 0.002 | 2b |
| CTR vs CA | 1-Phenyl-3-methyl-5-pyrazolone | 0.652 | -0.616 | 0.014 | 2b |
| CTR vs CA | Phosphoenolpyruvic acid | 0.649 | -0.625 | 0.002 | 2a |
| CTR vs CA | N-Iso valerylglycine | 0.647 | -0.627 | 0.002 | 1 |
| CTR vs CA | Malic acid | 0.647 | -0.628 | 0.033 | 1 |
| CTR vs CA | 4-Hydroxyproline | 0.633 | -0.659 | 7.740E-07 | 2a |
| CTR vs CA | N3,N4-Dimethyl-L-arginine | 0.632 | -0.663 | 4.210E-04 | 2b |
| CTR vs CA | N-Acetylneuraminic acid | 0.629 | -0.669 | 2.760E-04 | 2a |
| CTR vs CA | Acetylcholine | 0.623 | -0.683 | 3.090E-07 | 2b |
| CTR vs CA | Hypotaurine | 0.622 | -0.686 | 0.026 | 2a |
| CTR vs CA | Glycolic acid | 0.617 | -0.697 | 3.580E-06 | 2a |
| CTR vs CA | N6,N6,N6-Trimethyl-L-lysine | 0.616 | -0.698 | 9.620E-06 | 2b |
| CTR vs CA | N-Isobutyrylglycine | 0.605 | -0.725 | 1.340E-03 | 2b |
| CTR vs CA | 4-Acetamidophenol | 0.588 | -0.766 | 1.340E-03 | 2a |
| CTR vs CA | Citrulline | 0.588 | -0.766 | 2.580E-08 | 2a |
| CTR vs CA | 4-Hydroxyhippuric acid | 0.581 | -0.784 | 6.290E-04 | 1 |
| CTR vs CA | Phenol_1 | 0.578 | -0.790 | 3.410E-04 | 2b |
| CTR vs CA | 1-Methyl Adenosine | 0.577 | -0.794 | 4.210E-04 | 2a |
| CTR vs CA | 4,8-Dihydroxyquinoline-2-carboxylic acid | 0.576 | -0.796 | 1.110E-04 | 2a |
| CTR vs CA | Threonic acid | 0.574 | -0.801 | 4.210E-04 | 2a |
| CTR vs CA | 1H-indene-3-carboxamide | 0.568 | -0.816 | 0.006 | 2b |
| CTR vs CA | Pyridoxal | 0.554 | -0.853 | 1.310E-05 | 2a |
| CTR vs CA | Homoarginine | 0.533 | -0.909 | 2.580E-08 | 2a |
| CTR vs CA | Succinic acid | 0.524 | -0.931 | 0.037 | 1 |
| CTR vs CA | Methanesulfonic acid | 0.516 | -0.954 | 3.100E-05 | 2a |
| CTR vs CA | Serotonin | 0.511 | -0.970 | 0.004 | 1 |
| CTR vs CA | 7-Oxobenz[de]anthracene | 0.491 | -1.027 | 1.110E-03 | 2b |
| CTR vs CA | Trimethylamine N-Oxide | 0.489 | -1.033 | 0.002 | 2a |
| CTR vs CA | N-Acetyl-tyrosine | 0.479 | -1.062 | 7.650E-04 | 2a |
| CTR vs CA | Trigonelline | 0.464 | -1.107 | 7.650E-04 | 1 |
| CTR vs CA | Uracil | 0.463 | -1.110 | 0.008 | 2a |
| CTR vs CA | Xanthosine | 0.453 | -1.142 | 1.770E-04 | 2a |
| CTR vs CA | Guanidoacetic acid | 0.446 | -1.164 | 1.310E-05 | 1 |
| CTR vs CA | 3,4-Dihydroxybenzenesulfonic acid | 0.436 | -1.197 | 0.008 | 2b |
| CTR vs CA | 5-Hydroxyindole | 0.426 | -1.232 | 2.210E-04 | 2b |
| CTR vs CA | 2-Aminoadipic acid | 0.417 | -1.261 | 5.030E-06 | 2a |
| CTR vs CA | Acetyl-β-methylcholine_2 | 0.417 | -1.262 | 3.410E-04 | 2b |
| CTR vs CA | Catechol | 0.416 | -1.266 | 0.008 | 2b |
| CTR vs CA | Hexanoyl Glycine | 0.409 | -1.291 | 0.023 | 2a |
| CTR vs CA | Nicotinamide 1-oxide | 0.405 | -1.302 | 1.760E-05 | 2b |
| CTR vs CA | 3-Indoxylsulfuric acid | 0.404 | -1.307 | 2.760E-04 | 1 |
| CTR vs CA | 2-[(1S)-1-Hydroxyethyl]-4(1H)-quinazolinone | 0.401 | -1.320 | 0.002 | 2b |
| CTR vs CA | 3-(2-Hydroxyethyl)indole | 0.349 | -1.517 | 5.160E-04 | 2a |
| CTR vs CA | N-Acetyl-arginine | 0.338 | -1.566 | 3.100E-05 | 1 |
| CTR vs CA | Hippuric acid | 0.330 | -1.600 | 1.760E-05 | 1 |
| CTR vs CA | Ecgonine_2 | 0.318 | -1.651 | 1.310E-05 | 2b |
| CTR vs CA | Inosine | 0.268 | -1.902 | 2.500E-06 | 2a |
| CTR vs CA | Nitrosoheptamethyleneimine | 0.266 | -1.910 | 1.310E-05 | 2b |
| CTR vs CA | Thiamine | 0.231 | -2.117 | 3.100E-05 | 1 |
| CTR vs CA | 1,4a-Dimethyl-6-methylene-5-[2-(2-oxo-2,5-dihydro-3-furanyl)ethyl]decahydro-1-naphthalenecarboxylic acid | 0.214 | -2.223 | 2.580E-08 | 2b |
| CTR vs CA | 3-Aminosalicylic acid | 0.196 | -2.350 | 1.310E-05 | 2b |
| CTR vs CA | DL-Stachydrine_1 | 0.180 | -2.473 | 3.560E-05 | 2b |
| CTR vs CA | Glutathione oxidized | 0.119 | -3.076 | 1.760E-05 | 2a |
| CA vs VOR | SAICA-riboside | 2.343 | 1.228 | 1.730E-06 | 2a |
| CA vs VOR | Nitrosoheptamethyleneimine | 1.883 | 0.913 | 0.005 | 2b |
| CA vs VOR | Dehydronorketamine_1 | 1.560 | 0.642 | 0.033 | 2b |
| CA vs VOR | 3,5-Dimethoxybenzoic acid | 0.666 | -0.587 | 0.002 | 2b |
| CA vs VOR | Cytosine | 0.607 | -0.720 | 0.006 | 1 |
| CA vs VOR | 3-(propan-2-yl)-octahydropyrrolo[1,2-a]pyrazine-1,4-dione_1 | 0.506 | -0.982 | 0.023 | 2b |
| CA vs VOR | 2-[(1S)-1-Hydroxyethyl]-4(1H)-quinazolinone | 0.430 | -1.217 | 0.023 | 2b |
| CA vs VOR | (8aR,12S,12aR)-12-Hydroxy-4-methyl-4,5,6,7,8,8a,12,12a-octahydro-2H-3-benzoxecine-2,9(1H)-dione | 0.402 | -1.315 | 3.580E-06 | 2b |
| CA vs VOR | Glycocholic acid | 0.393 | -1.347 | 0.033 | 1 |
| CA vs VOR | Hyocholic acid | 0.387 | -1.368 | 0.004 | 2a |
| CA vs VOR | Cholic acid | 0.378 | -1.402 | 0.026 | 1 |
| CA vs VOR | 5,6-dimethyl-4-oxo-4H-pyran-2-carboxylic acid | 0.361 | -1.468 | 1.410E-04 | 2b |
| CA vs VOR | 3-(propan-2-yl)-octahydropyrrolo[1,2-a]pyrazine-1,4-dione_2 | 0.356 | -1.490 | 6.290E-04 | 2b |
| CA vs VOR | 2-Hydroxyphenylalanine | 0.260 | -1.943 | 5.160E-08 | 2b |
| CA vs WT | Thiamine | 2.340 | 1.227 | 0.004 | 1 |
| CA vs WT | Aspartic acid | 1.671 | 0.741 | 0.027 | 2a |
| CA vs WT | 3-Methylhistidine | 1.535 | 0.618 | 3.720E-05 | 1 |
| CA vs WT | Creatine | 1.504 | 0.589 | 0.002 | 1 |
| CA vs WT | Isovalerylcarnitine | 0.663 | -0.592 | 9.870E-04 | 1 |
| CA vs WT | N-Acetyl-ornithine | 0.662 | -0.594 | 1.320E-03 | 1 |
| CA vs WT | Pyridoxine | 0.652 | -0.616 | 0.003 | 2a |
| CA vs WT | 2,3,4-Trihydroxybenzoic acid | 0.648 | -0.626 | 0.023 | 2a |
| CA vs WT | β-D-Glucopyranuronic acid | 0.633 | -0.659 | 0.040 | 2b |
| CA vs WT | Prolylglycine | 0.628 | -0.672 | 2.270E-06 | 2b |
| CA vs WT | Acetyl-β-methylcholine_1 | 0.623 | -0.682 | 0.012 | 2b |
| CA vs WT | 1,4a-Dimethyl-6-methylene-5-[2-(2-oxo-2,5-dihydro-3-furanyl)ethyl]decahydro-1-naphthalenecarboxylic acid | 0.606 | -0.724 | 1.010E-05 | 2b |
| CA vs WT | Homocitrulline | 0.605 | -0.726 | 3.720E-05 | 2a |
| CA vs WT | N-Acetyl-lysine | 0.603 | -0.731 | 8.510E-04 | 1 |
| CA vs WT | Ferulic acid | 0.602 | -0.731 | 0.007 | 2a |
| CA vs WT | 3-(1-hydroxyethyl)-2,3,6,7,8,8a-hexahydropyrrolo[1,2-a]pyrazine-1,4-dione | 0.597 | -0.744 | 0.019 | 2b |
| CA vs WT | Dehydronorketamine_3 | 0.590 | -0.761 | 0.015 | 2b |
| CA vs WT | 2-Hydroxyisocaproic acid | 0.587 | -0.769 | 1.980E-05 | 2a |
| CA vs WT | Methionine sulfoxide | 0.581 | -0.784 | 0.009 | 2a |
| CA vs WT | Indole-3-acetic acid | 0.579 | -0.790 | 2.830E-04 | 2a |
| CA vs WT | Acetyl-β-methylcholine_2 | 0.565 | -0.824 | 0.005 | 2b |
| CA vs WT | trans-3-Hydroxycinnamic acid | 0.533 | -0.907 | 0.002 | 2a |
| CA vs WT | SAICA-riboside | 0.523 | -0.936 | 0.005 | 2a |
| CA vs WT | [(1S,4S,6S)-6-Isopropyl-3-methyl-4-{2-oxo-2-[4-(2-pyrimidinyl)-1-piperazinyl]ethyl}-2-cyclohexen-1-yl]acetonitrile | 0.503 | -0.991 | 1.140E-03 | 2b |
| CA vs WT | 4-Coumaric acid | 0.497 | -1.007 | 0.024 | 2a |
| CA vs WT | Cytosine | 0.494 | -1.019 | 6.290E-04 | 1 |
| CA vs WT | 3-(propan-2-yl)-octahydropyrrolo[1,2-a]pyrazine-1,4-dione_2 | 0.492 | -1.024 | 0.002 | 2b |
| CA vs WT | 5-Methoxytryptophan | 0.488 | -1.035 | 0.014 | 2a |
| CA vs WT | N-Acetyl-5-hydroxytryptamine | 0.458 | -1.127 | 2.830E-04 | 2a |
| CA vs WT | 5-Methylcytosine | 0.437 | -1.193 | 1.270E-05 | 2a |
| CA vs WT | DL-Stachydrine_2 | 0.431 | -1.213 | 1.030E-04 | 2b |
| CA vs WT | 10-HDA | 0.424 | -1.237 | 0.002 | 2b |
| CA vs WT | 4-cyclopropyl-6-methoxy-1,3,5-triazin-2-amine | 0.411 | -1.282 | 0.012 | 2b |
| CA vs WT | Hippuric acid | 0.379 | -1.398 | 0.003 | 1 |
| CA vs WT | Trigonelline | 0.335 | -1.578 | 0.003 | 1 |
| CA vs WT | Catechol | 0.334 | -1.581 | 0.002 | 2b |
| CA vs WT | 4-Hydroxybenzaldehyde | 0.332 | -1.589 | 3.740E-05 | 2a |
| CA vs WT | (R)-Equol | 0.320 | -1.643 | 8.000E-06 | 2b |
| CA vs WT | 3-(2-Hydroxyethyl)indole | 0.315 | -1.666 | 0.033 | 2a |
| CA vs WT | Indole-3-acrylic acid | 0.276 | -1.856 | 1.310E-06 | 2a |
| CA vs WT | (8aR,12S,12aR)-12-Hydroxy-4-methyl-4,5,6,7,8,8a,12,12a-octahydro-2H-3-benzoxecine-2,9(1H)-dione | 0.269 | -1.894 | 9.630E-08 | 2b |
| CA vs WT | 2-Hydroxyphenylalanine | 0.254 | -1.975 | 9.810E-07 | 2b |
| CA vs WT | 3,4-Dihydroxybenzenesulfonic acid | 0.250 | -2.001 | 0.002 | 2b |
| CA vs WT | Hyocholic acid | 0.230 | -2.120 | 0.017 | 2a |
| CA vs WT | 3-(propan-2-yl)-octahydropyrrolo[1,2-a]pyrazine-1,4-dione_1 | 0.223 | -2.164 | 3.830E-06 | 2b |
| CA vs WT | Indole-3-propionic acid | 0.216 | -2.213 | 8.000E-06 | 2a |
| CA vs WT | a/b/ω-Muricholic acid | 0.139 | -2.846 | 8.510E-04 | 2a |
| CA vs WT | Cholic acid | 0.124 | -3.014 | 0.010 | 1 |
| CA vs PptA | cis-Aconitic acid | 7.053 | 2.818 | 1.270E-05 | 2a |
| CA vs PptA | Succinic acid | 5.333 | 2.415 | 0.006 | 1 |
| CA vs PptA | 3-(1-hydroxyethyl)-2,3,6,7,8,8a-hexahydropyrrolo[1,2-a]pyrazine-1,4-dione | 5.095 | 2.349 | 2.270E-06 | 2b |
| CA vs PptA | Thiamine | 4.983 | 2.317 | 2.000E-07 | 1 |
| CA vs PptA | 7-Oxobenz[de]anthracene | 4.687 | 2.229 | 4.930E-06 | 2b |
| CA vs PptA | 4-Coumaric acid | 4.642 | 2.215 | 2.270E-06 | 2a |
| CA vs PptA | Mucic acid | 4.452 | 2.154 | 8.160E-05 | 2a |
| CA vs PptA | 4-cyclopropyl-6-methoxy-1,3,5-triazin-2-amine | 4.185 | 2.065 | 6.290E-04 | 2b |
| CA vs PptA | 3-(propan-2-yl)-octahydropyrrolo[1,2-a]pyrazine-1,4-dione_2 | 3.570 | 1.836 | 3.830E-06 | 2b |
| CA vs PptA | Daidzein | 3.375 | 1.755 | 5.390E-04 | 2a |
| CA vs PptA | 4,8-Dihydroxyquinoline-2-carboxylic acid | 3.261 | 1.706 | 3.830E-06 | 2a |
| CA vs PptA | 5-Methoxytryptophan | 3.161 | 1.660 | 1.980E-05 | 2a |
| CA vs PptA | Genistein | 3.138 | 1.650 | 9.830E-05 | 2a |
| CA vs PptA | 3-(propan-2-yl)-octahydropyrrolo[1,2-a]pyrazine-1,4-dione_1 | 3.038 | 1.603 | 2.960E-06 | 2b |
| CA vs PptA | DL-Stachydrine_1 | 2.864 | 1.518 | 4.280E-04 | 2b |
| CA vs PptA | Pyridoxal | 2.799 | 1.485 | 1.440E-09 | 2a |
| CA vs PptA | Inosine | 2.773 | 1.471 | 0.020 | 2a |
| CA vs PptA | DL-Stachydrine_2 | 2.729 | 1.448 | 3.030E-05 | 2b |
| CA vs PptA | Trigonelline | 2.691 | 1.428 | 1.980E-05 | 1 |
| CA vs PptA | Uracil | 2.453 | 1.295 | 0.047 | 2a |
| CA vs PptA | 3,4-Dihydroxybenzenesulfonic acid | 2.408 | 1.268 | 1.320E-03 | 2b |
| CA vs PptA | Catechol | 2.398 | 1.262 | 0.002 | 2b |
| CA vs PptA | α-aminobutyric acid/γ-Aminobutyric acid | 2.393 | 1.259 | 4.310E-08 | 2a |
| CA vs PptA | Biotin | 2.361 | 1.240 | 8.000E-06 | 2a |
| CA vs PptA | N-Acetyl-glutamic acid | 2.334 | 1.223 | 9.830E-05 | 2a |
| CA vs PptA | Ferulic acid | 2.317 | 1.212 | 7.330E-04 | 2a |
| CA vs PptA | Diaminopimelic acid | 2.279 | 1.189 | 1.010E-05 | 2a |
| CA vs PptA | Acetyl-β-methylcholine_2 | 2.278 | 1.188 | 6.300E-06 | 2b |
| CA vs PptA | N-Acetyl-tyrosine | 2.200 | 1.138 | 2.270E-06 | 2a |
| CA vs PptA | Sugar alcochol (C5) | 2.183 | 1.126 | 1.980E-05 | 2a |
| CA vs PptA | Glutathione oxidized | 2.158 | 1.110 | 0.005 | 2a |
| CA vs PptA | 2,3,4-Trihydroxybenzoic acid | 2.137 | 1.096 | 0.007 | 2a |
| CA vs PptA | trans-3-Hydroxycinnamic acid | 2.105 | 1.074 | 1.140E-03 | 2a |
| CA vs PptA | Kynurenic acid | 2.092 | 1.065 | 5.550E-05 | 2a |
| CA vs PptA | Methionine sulfoxide | 2.076 | 1.054 | 2.960E-06 | 2a |
| CA vs PptA | Gluconic acid | 1.900 | 0.926 | 4.610E-04 | 1 |
| CA vs PptA | Xanthosine | 1.833 | 0.874 | 0.030 | 2a |
| CA vs PptA | Methanesulfonic acid | 1.777 | 0.829 | 8.000E-06 | 2a |
| CA vs PptA | Malic acid | 1.761 | 0.816 | 8.510E-04 | 1 |
| CA vs PptA | 5-Hydroxyindole | 1.756 | 0.813 | 1.690E-04 | 2b |
| CA vs PptA | 3-Indoxylsulfuric acid | 1.739 | 0.798 | 1.180E-04 | 1 |
| CA vs PptA | δ-Gluconolactone | 1.678 | 0.747 | 7.330E-04 | 2a |
| CA vs PptA | Glyoxylic acid | 1.667 | 0.737 | 8.510E-04 | 2a |
| CA vs PptA | Pyridoxine | 1.666 | 0.737 | 0.006 | 2a |
| CA vs PptA | Betaine | 1.666 | 0.736 | 2.000E-07 | 1 |
| CA vs PptA | Proline | 1.624 | 0.699 | 6.740E-05 | 1 |
| CA vs PptA | N-Acetyl-arginine | 1.622 | 0.698 | 1.320E-03 | 1 |
| CA vs PptA | Hippuric acid | 1.620 | 0.696 | 0.011 | 1 |
| CA vs PptA | Uridine | 1.600 | 0.678 | 0.043 | 1 |
| CA vs PptA | DL-4-Hydroxyphenyllactic acid | 1.545 | 0.627 | 0.004 | 2b |
| CA vs PptA | Acetanilide | 1.543 | 0.626 | 3.830E-06 | 2b |
| CA vs PptA | Fumaric acid | 1.539 | 0.622 | 2.830E-04 | 1 |
| CA vs PptA | Tyrosine | 1.534 | 0.618 | 1.980E-05 | 1 |
| CA vs PptA | 3-Ureidopropionic acid | 1.527 | 0.611 | 0.012 | 2a |
| CA vs PptA | N-Acetyl-lysine | 1.507 | 0.592 | 5.390E-04 | 1 |
| CA vs PptA | Coumarin | 1.500 | 0.585 | 1.010E-05 | 2b |
| CA vs PptA | Isovalerylcarnitine | 0.662 | -0.595 | 1.690E-04 | 1 |
| CA vs PptA | 3-Methyl-2-oxovaleric acid | 0.649 | -0.624 | 3.920E-04 | 2a |
| CA vs PptA | Dehydronorketamine_1 | 0.625 | -0.677 | 0.015 | 2b |
| CA vs PptA | Tiglic acid | 0.617 | -0.697 | 0.003 | 2b |
| CA vs PptA | 4-Hydroxy xylazine | 0.608 | -0.718 | 0.044 | 2b |
| CA vs PptA | Norketamine_1 | 0.597 | -0.743 | 0.015 | 2b |
| CA vs PptA | Pyrene | 0.596 | -0.747 | 0.001 | 2b |
| CA vs PptA | [(1S,4S,6S)-6-Isopropyl-3-methyl-4-{2-oxo-2-[4-(2-pyrimidinyl)-1-piperazinyl]ethyl}-2-cyclohexen-1-yl]acetonitrile | 0.590 | -0.760 | 8.510E-04 | 2b |
| CA vs PptA | SAICA-riboside | 0.586 | -0.771 | 0.007 | 2a |
| CA vs PptA | Acetyl-β-methylcholine_1 | 0.574 | -0.801 | 0.002 | 2b |
| CA vs PptA | Citric acid | 0.550 | -0.862 | 1.420E-04 | 1 |
| CA vs PptA | N-Acetyl-5-hydroxytryptamine | 0.538 | -0.894 | 0.033 | 2a |
| CA vs PptA | Crotonic acid_2 | 0.525 | -0.930 | 8.160E-05 | 2b |
| CA vs PptA | Rexamino | 0.511 | -0.969 | 0.024 | 2b |
| CA vs PptA | 10-HDA | 0.509 | -0.975 | 0.006 | 2b |
| CA vs PptA | Cortisone | 0.503 | -0.992 | 0.027 | 1 |
| CA vs PptA | Butyrylcarnitine | 0.500 | -0.999 | 3.910E-07 | 2a |
| CA vs PptA | 11-α-Hydroxy-17-methyltestosterone | 0.487 | -1.039 | 2.010E-04 | 2b |
| CA vs PptA | Indole-3-acetic acid | 0.480 | -1.058 | 6.740E-05 | 2a |
| CA vs PptA | N8-Acetylspermidine | 0.480 | -1.059 | 1.420E-04 | 2a |
| CA vs PptA | 20 β-Dihydroprednisolone | 0.424 | -1.237 | 0.010 | 2b |
| CA vs PptA | Dehydronorketamine_3 | 0.398 | -1.329 | 9.870E-04 | 2b |
| CA vs PptA | Cytosine | 0.345 | -1.534 | 9.830E-05 | 1 |
| CA vs PptA | N-Acetyl-cysteine | 0.343 | -1.542 | 0.004 | 1 |
| CA vs PptA | Octanoylcarnitine | 0.340 | -1.556 | 2.270E-06 | 2a |
| CA vs PptA | 3-Hydroxy-2-methyl-4-pyrone | 0.333 | -1.586 | 1.980E-05 | 2a |
| CA vs PptA | 3-Hydroxybutyric acid | 0.330 | -1.598 | 8.160E-05 | 1 |
| CA vs PptA | Glycocholic acid | 0.285 | -1.809 | 0.010 | 1 |
| CA vs PptA | (-)-Camphanic acid | 0.285 | -1.812 | 8.000E-06 | 2b |
| CA vs PptA | 2-Hydroxyoctanoic acid | 0.255 | -1.970 | 2.270E-06 | 1 |
| CA vs PptA | Hyocholic acid | 0.215 | -2.218 | 1.180E-04 | 2a |
| CA vs PptA | Decanoylcarnitine | 0.118 | -3.088 | 5.750E-09 | 1 |
| CA vs PptA | a/b/ω-Muricholic acid | 0.074 | -3.764 | 3.330E-04 | 2a |
| CA vs PptA | Cholic acid | 0.061 | -4.040 | 2.010E-04 | 1 |
| WT vs Tx | DL-Stachydrine_1 | 6.876 | 2.782 | 3.810E-06 | 2b |
| WT vs Tx | 3,4-Dihydroxybenzenesulfonic acid | 6.338 | 2.664 | 3.980E-05 | 2b |
| WT vs Tx | Catechol | 4.700 | 2.233 | 4.700E-05 | 2b |
| WT vs Tx | 3-(propan-2-yl)-octahydropyrrolo[1,2-a]pyrazine-1,4-dione_1 | 4.528 | 2.179 | 2.790E-06 | 2b |
| WT vs Tx | Trigonelline | 4.405 | 2.139 | 3.400E-07 | 1 |
| WT vs Tx | Daidzein | 3.610 | 1.852 | 0.003 | 2a |
| WT vs Tx | 4-Coumaric acid | 3.596 | 1.846 | 1.670E-04 | 2a |
| WT vs Tx | 3-(2-Hydroxyethyl)indole | 3.451 | 1.787 | 0.024 | 2a |
| WT vs Tx | Hippuric acid | 3.260 | 1.705 | 3.900E-04 | 1 |
| WT vs Tx | 5-Methoxytryptophan | 3.249 | 1.700 | 2.250E-06 | 2a |
| WT vs Tx | SAICA-riboside | 3.238 | 1.695 | 2.820E-05 | 2a |
| WT vs Tx | Indole-3-propionic acid | 3.170 | 1.665 | 4.700E-05 | 2a |
| WT vs Tx | cis-Aconitic acid | 2.997 | 1.583 | 0.020 | 2a |
| WT vs Tx | trans-3-Hydroxycinnamic acid | 2.929 | 1.550 | 2.420E-05 | 2a |
| WT vs Tx | 4-Guanidinobutyric acid | 2.741 | 1.455 | 0.004 | 1 |
| WT vs Tx | 7-Oxobenz[de]anthracene | 2.677 | 1.421 | 1.050E-04 | 2b |
| WT vs Tx | DL-Stachydrine_2 | 2.648 | 1.405 | 2.070E-05 | 2b |
| WT vs Tx | 4-Hydroxybenzaldehyde | 2.579 | 1.367 | 1.350E-05 | 2a |
| WT vs Tx | Ferulic acid | 2.553 | 1.352 | 1.450E-04 | 2a |
| WT vs Tx | (R)-Equol | 2.407 | 1.267 | 9.120E-07 | 2b |
| WT vs Tx | 3-(1-hydroxyethyl)-2,3,6,7,8,8a-hexahydropyrrolo[1,2-a]pyrazine-1,4-dione | 2.359 | 1.238 | 9.410E-06 | 2b |
| WT vs Tx | Genistein | 2.304 | 1.204 | 0.013 | 2a |
| WT vs Tx | Acetyl-β-methylcholine_2 | 2.297 | 1.200 | 2.790E-06 | 2b |
| WT vs Tx | Sugar alcochol (C5) | 2.162 | 1.113 | 1.370E-05 | 2a |
| WT vs Tx | Methionine sulfoxide | 2.104 | 1.073 | 7.570E-04 | 2a |
| WT vs Tx | Diaminopimelic acid | 2.088 | 1.062 | 5.210E-06 | 2a |
| WT vs Tx | Dehydronorketamine_1 | 1.830 | 0.872 | 0.008 | 2b |
| WT vs Tx | N-Acetyl-lysine | 1.824 | 0.867 | 9.750E-04 | 1 |
| WT vs Tx | Homocitrulline | 1.765 | 0.820 | 4.690E-08 | 2a |
| WT vs Tx | 2,3,4-Trihydroxybenzoic acid | 1.709 | 0.773 | 0.010 | 2a |
| WT vs Tx | Indole-3-acrylic acid | 1.708 | 0.773 | 0.004 | 2a |
| WT vs Tx | (8aR,12S,12aR)-12-Hydroxy-4-methyl-4,5,6,7,8,8a,12,12a-octahydro-2H-3-benzoxecine-2,9(1H)-dione | 1.703 | 0.768 | 1.980E-05 | 2b |
| WT vs Tx | Prolylglycine | 1.700 | 0.766 | 2.010E-07 | 2b |
| WT vs Tx | N-Acetyl-tyrosine | 1.681 | 0.749 | 0.031 | 2a |
| WT vs Tx | 5-Methylcytosine | 1.649 | 0.721 | 2.560E-04 | 2a |
| WT vs Tx | Glucuronic acid | 1.601 | 0.679 | 0.002 | 2a |
| WT vs Tx | α-aminobutyric acid/γ-Aminobutyric acid | 1.588 | 0.668 | 2.450E-08 | 2a |
| WT vs Tx | Pyridoxal | 1.534 | 0.617 | 0.015 | 2a |
| WT vs Tx | N-Acetyl-ornithine | 1.513 | 0.598 | 2.820E-05 | 1 |
| WT vs Tx | 3,4-Dimethoxyphenylacetic acid | 0.664 | -0.591 | 0.026 | 1 |
| WT vs Tx | AcetylCarnitine | 0.619 | -0.692 | 1.920E-04 | 1 |
| WT vs Tx | Isobutyric acid_1 | 0.616 | -0.699 | 0.033 | 2b |
| WT vs Tx | Ethylmalonic acid | 0.605 | -0.726 | 0.033 | 1 |
| WT vs Tx | 3-Aminosalicylic acid | 0.588 | -0.766 | 0.008 | 2b |
| WT vs Tx | Triisopropanolamine | 0.579 | -0.788 | 0.002 | 2b |
| WT vs Tx | Octanoylcarnitine | 0.570 | -0.812 | 0.049 | 2a |
| WT vs Tx | 20 β-Dihydroprednisolone | 0.559 | -0.840 | 0.010 | 2b |
| WT vs Tx | 3,5-Dimethoxybenzoic acid | 0.546 | -0.873 | 5.840E-04 | 2b |
| WT vs Tx | Citric acid | 0.540 | -0.888 | 0.018 | 1 |
| WT vs Tx | 5-Methoxysalicylic acid | 0.537 | -0.897 | 0.017 | 2a |
| WT vs Tx | 3-Hydroxy-2-methyl-4-pyrone | 0.528 | -0.922 | 0.007 | 2a |
| WT vs Tx | Decanoylcarnitine | 0.484 | -1.046 | 4.470E-04 | 1 |
| WT vs Tx | 4-Hydroxy xylazine | 0.482 | -1.054 | 1.100E-03 | 2b |
| WT vs Tx | 5,6-dimethyl-4-oxo-4H-pyran-2-carboxylic acid | 0.471 | -1.086 | 3.900E-04 | 2b |
| WT vs Tx | Crotonic acid_2 | 0.458 | -1.127 | 3.400E-04 | 2b |
| WT vs Tx | N6,N6,N6-Trimethyl-L-lysine | 0.450 | -1.152 | 8.600E-04 | 2b |
| WT vs Tx | 2-Hydroxyoctanoic acid | 0.447 | -1.161 | 0.002 | 1 |
| WT vs Tx | (-)-Camphanic acid | 0.420 | -1.252 | 8.600E-04 | 2b |
| WT vs Tx | 11- α-Hydroxy-17-methyltestosterone | 0.387 | -1.369 | 0.004 | 2b |
| WT vs Tx | 3-Hydroxybutyric acid | 0.286 | -1.807 | 1.100E-03 | 1 |
